# Linking enzyme expression to metabolic flux

**DOI:** 10.1101/2022.11.17.516982

**Authors:** Xuhang Li, Albertha J.M. Walhout, L. Safak Yilmaz

## Abstract

Metabolic reaction flux is regulated in response to nutritional, environmental or pathological conditions by changes in either metabolite or metabolic enzyme levels. Previous studies proposed that flux is predominately regulated by metabolite, rather than enzyme, levels. However, the extent to which changes in enzyme levels affect flux throughout the metabolic network remains unclear. Here, we combine available yeast enzyme level, flux data, and metabolic network modeling to demonstrate three paradigms by which enzyme levels are broadly associated with flux: cognate reaction, pathway-level coordination, and flux coupling. We find that the architecture of the metabolic network enables the reach of influence for most enzymes. We implemented enzyme reach as a novel parameter in an enhanced flux potential analysis algorithm, which predicts relative flux levels under different conditions from variations in enzyme expression. This algorithm was tested in yeast and humans. Our study suggests that metabolic network architecture facilitates a broad physiological impact of changes in enzyme levels and may form a foundation for using enzyme expression data for a variety of systems, and eventually, individual cells.

## Introduction

Metabolism is highly dynamic. Metabolic flux, or the rate of metabolic conversion, changes in response to nutritional, environmental, and pathogenic perturbations. Flux can be regulated by multiple mechanisms: metabolite concentrations can directly affect reaction rate; allosteric regulators and covalent modifications can affect the activity of metabolic enzymes; and metabolic enzyme levels can be regulated by changes in gene or protein expression (Fell, 1997). The last mechanism can serve two purposes: the first is innate and is to establish enzyme levels in different cell types or during different developmental stages. The second is in response to exogenous conditions, such as changes in diet or exposure to specific compounds. The first is what we refer to as metabolic wiring, while the second can be viewed as rewiring. Overall, the relationships between enzyme levels, wiring and rewiring of metabolic flux at a systems, or network, level remain elusive (Kochanowski *et al*, 2013).

To fully understand the role of enzyme expression in metabolic (re)wiring, two critical elements are required: a comprehensive dataset encompassing flux and enzyme abundance measurements that span a significant portion of a metabolic network, and methodologies capable of mathematically linking enzyme levels to flux within this network. The acquisition of fluxomic data covering a substantial number of reactions can be challenging. Conventional approaches that determine flux through full isotope balancing are often limited to core carbon metabolic pathways (Gopalakrishnan & Maranas, 2015; Kharchenko *et al*, 2005; Zamboni *et al*, 2009). Alternatively, network-level flux distributions can be predicted using Flux Balance Analysis (FBA) with a wide range of measured rates as constraints (Antoniewicz, 2015; Hackett *et al*, 2016)). However, the number of reactions for which the solution space can be sufficiently constrained, i.e., where flux can be determined, largely depends on the comprehensiveness of the measured rates. Consequently, to the best of our knowledge, only a handful of studies to date have achieved network-level flux determination (Hackett *et al*., 2016; Kochanowski *et al*, 2021; Lahtvee *et al*, 2017).

Studies that simultaneously measured flux and enzyme levels generally suggest that flux is primarily controlled by metabolite-, rather than enzyme levels (Chubukov *et al*, 2013; Daran-Lapujade *et al*, 2004; Daran-Lapujade *et al*, 2007; Hackett *et al*., 2016; Lahtvee *et al*., 2017; Yu *et al*, 2020). However, these studies largely employ data integration methodologies that focus on regulation at the individual reaction level. In contrast, metabolic control analysis (MCA) theory posits that changes in an enzyme’s activity can propagate influences throughout the network and that multiple enzymes jointly control a pathway’s flux (Kacser & Burns, 1973). This makes intuitive sense since, at steady state, the producing and consuming fluxes of the same metabolite need to be balanced, resulting in flux coupling of related reactions. Therefore, metabolic (re)wiring by gene expression changes may affect not only cognate reactions, but also have a larger reach through the network.

Methods for network-level integration have been formulated to embed enzyme expression data in a metabolic network context, aiming to infer metabolic flux (Machado & Herrgard, 2014; Opdam *et al*, 2017). However, the extent to which flux can be quantitatively associated with network-integrated enzyme levels remains unclear. Most of the past studies focused on reconstructing context-specific metabolic networks to predict metabolic wiring in tissues and cells of multicellular organisms (Li *et al*, 2022). These analyses are limited by qualitative flux predictions and the lack of experimental flux data for validation (Agren *et al*, 2012; Becker & Palsson, 2008; Jensen & Papin, 2011; Jerby *et al*, 2010; Vlassis *et al*, 2014). Several studies attempted to predict absolute flux levels using enzyme levels as additional constraints in the framework of FBA (Colijn *et al*, 2009; Lee *et al*, 2012; O’Brien *et al*, 2013; Salvy & Hatzimanikatis, 2020; Sanchez *et al*, 2017). However, the integration of enzyme levels usually does not enhance flux prediction when benchmarked against experimental fluxes measured in microorganisms (Machado & Herrgard, 2014). More recent studies turned to network-integration analysis of relative changes in enzyme levels and flux across different conditions (Kim & Reed, 2012; Pandey *et al*, 2019; Pusa *et al*, 2020; Ravi & Gunawan, 2021; Zhu *et al*, 2017) (Gavai *et al*, 2015), but these tools have not been used to dissect flux regulation at a systems level. We recently developed Flux Potential Analysis (FPA), which integrates the relative enzyme levels under different conditions of not only the cognate enzyme for a reaction of interest but also the enzymes of nearby reactions within the same network neighborhood (Yilmaz *et al*, 2020). FPA provides an opportunity to computationally interrogate the role of changes in enzyme expression in flux (re)wiring within the context of a metabolic network.

In this study, we investigated the influence of enzyme level variations within the metabolic network of yeast, using published network-level flux estimates and proteomic data from 25 distinct growth conditions. We first developed an enhanced FPA (eFPA) algorithm to integrate expression levels within a more precisely defined network neighborhood that we refer to as relevant pathway(s). Using eFPA, we found that changes in flux are largely associated with the coordinated regulation of enzyme expression in the network, particularly within the immediate (local) pathway. This suggests that the impact of enzyme expression is considerably greater than previously assumed based on reaction-level analyses. We also applied the eFPA algorithm, calibrated with yeast data, to predict relative levels of flux in human tissues. This analysis not only validated our findings in a wholly different system, but also revealed interesting insights on tissue metabolic functions.

## Results

To systematically evaluate the role of enzyme regulation in flux (re)wiring, we required a dataset that *(i)* had both flux and enzyme expression data acquired from the same samples; *(ii)* provided accurate flux values spread across a metabolic network, not just confined to core carbon metabolism; and *(iii)* involved multiple conditions to ensure a statistically meaningful analysis. Our literature survey led us to conclude that only the yeast dataset from Hackett et al. (Hackett *et al*., 2016) fulfilled these requirements (Table EV1). In this dataset, flux estimates and enzyme expression measurements are available for 156 metabolic reactions across 25 conditions in which different nutrients (glucose, leucine, uracil, phosphate, nitrogen) were limited and the growth rate was titrated (Tables EV2-EV4).

In order to compare enzyme expression with flux, it is essential that both variables – enzyme levels and flux – are contextualized appropriately. In yeast cultures, both protein abundance and flux have been shown to scale with specific growth rate (Hackett *et al*., 2016; Xia *et al*, 2022), which confounds the interpretation of a direct comparison. In Hackett et al, the proteomic data reflects enzyme abundance in terms of its proportion to the total protein (Hackett *et al*., 2016), making it intrinsically adjusted for growth variations. To ensure a meaningful comparison with flux, we adjusted the flux data using the corresponding growth rates. These adjusted flux values are used throughout this study (referred to as flux, see Appendix Supplementary Methods for details).

### Metabolic enzyme levels influence flux of non-cognate reactions

We first identified metabolic reactions for which flux correlates with the level of the corresponding enzyme(s) (Fig. 1a). A total of 46 of the 156 reactions (29%) showed a significant positive correlation between flux and enzyme levels (False Discovery Rate (FDR) < 0.05). Hereafter these reactions are referred to as *correlated reactions* (Fig. 1b, c, Appendix Fig. S1 and Table EV2). Similar to observations in bacteria (Chubukov *et al*., 2013), most central carbon reactions in yeast are not correlated, while correlated reactions mainly occur in amino acid and nucleic acid metabolism (Fig. 1d, Fig. EV1a-d). These findings agree with a recent study on proteome allocations which suggested that enzyme usage significantly impact flux in amino acid biosynthesis (Xia *et al*., 2022). Fluxes in arginine, phenylalanine, tryptophan and tyrosine biosynthesis are correlated, as found previously (Fig. 1d) (Lahtvee *et al*., 2017; Moxley *et al*, 2009). Cumulatively, our correlation analysis reaffirms the well-documented influence of enzyme expression on flux regulation at the individual reaction level. Interestingly, some metabolic pathways have multiple correlated reactions, while others have only one or a few. For instance, all seven reactions in arginine biosynthesis are correlated, whereas in threonine, methionine, and cysteine metabolism, only the final reaction is correlated (Fig. EV1b, d).

**Fig. 1:**
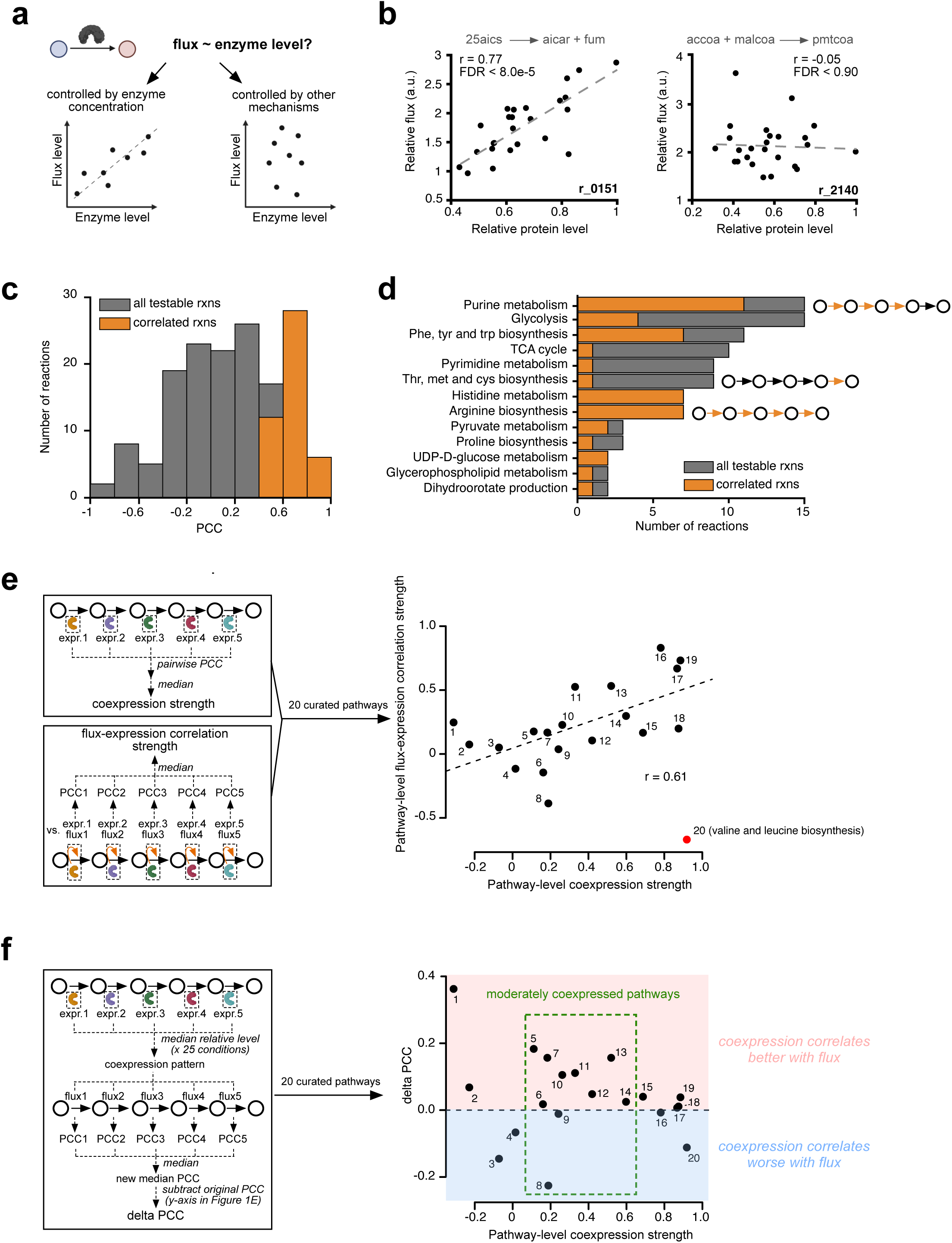
Pathway-level coordinated regulation of enzyme levels dictates metabolic flux in yeast. **a**, Models for quantitative relations between enzyme levels and flux. **b**, Representative examples for flux-enzyme level correlation. Each datapoint represents a measurement (25 total) of relative flux and enzyme levels for the indicated reaction. The metabolite abbreviations are defined by BiGG model nomenclature(King *et al*, 2016). a.u., arbitrary units. **c**, Pearson Correlation Coefficient (PCC) distribution for the 156 reactions for which flux and enzyme levels were both available. Reactions with significant positive correlation (FDR < 0.05, PCC > 0) are indicated in orange. **d**, Metabolic pathway annotations of the 156 reactions. **e**, Correlation between pathway-level enzyme coexpression and flux-expression concordance. Pathway information corresponding to the number indices is provided in Fig. EV1e. The trend line was fitted without the outlier (red dot). The correlation coefficient (r) of the trend line is shown in the plot. **f**, Comparing flux concordance with individual and pathway-level enzyme expression.

The yeast experiments were performed at steady state, and, therefore, the flux in reactions of the same linear pathway is coordinated (Hackett *et al*., 2016). Further, enzymes that catalyze reactions in the same pathway are often coordinately activated or repressed, thereby enabling network rewiring (Bulcha *et al*, 2019; Giese *et al*, 2020; Kochanowski *et al*., 2021; Nanda *et al*, 2023; Watson *et al*, 2016). Therefore, we next asked whether pathways with a greater degree of enzyme coexpression exhibit an overall better correlation between enzyme levels and flux. We manually defined 20 metabolic pathways that consist of at least three connected reactions for which flux estimations were available, and for which the enzyme levels corresponding to at least two reactions were available (Fig. EV1e and Table EV2). We found that, except for valine and leucine biosynthesis, there was a significant correlation between the strength of pathway-level enzyme coexpression and the strength of flux-expression correlation (r=0.61, p<0.006) (Fig. 1e). The exception of leucine biosynthesis pathway is likely due to the use of a leucine auxotrophic strain (MAT**a** *leu2Δ1*)(Hackett *et al*., 2016). The correlation indicates that the more different enzymes in a pathway covary in expression under different conditions, the more the flux through the entire pathway correlates with changes in enzyme levels. Based on this observation, we next asked if pathway-level coexpression better predicts flux variations compared to changes in individual enzymes. We defined pathway-level coexpression as a vector of 25 elements that corresponds to the 25 conditions, with each element containing the median of relative enzyme levels of all enzymes in the pathway in that condition. Indeed, enzyme coexpression correlates better with pathway flux than the enzyme levels for individual reactions, particularly for pathways that are moderately coexpressed (Fig. 1f). Therefore, we conclude that pathway-level regulation of enzyme expression better predicts flux changes compared with changes in enzyme levels of individual reactions. This observation supports the idea that changes in enzyme levels not only affect cognate reactions, but have a reach by which they affect other, non-cognate, reactions in the same pathway as well.

### Enzyme reach affects local pathway flux

Which non-cognate reactions are affected by changes in the level of individual enzymes, or, in other words, what is the extent of “enzyme reach”? We hypothesized that enzyme reach is determined by the network architecture. For instance, the reach is likely to extend along reactions that have a linear connection to the cognate reaction, especially since flux is coupled in linear pathways. Such linear connections are reflected in the network architecture and can be quantified by the network connectivity degree of metabolites (Fig. 2a). While a degree of two indicates strict linearity, low degrees may generally suggest higher likelihood of linear connections in actual flux. To test whether the metabolite degree is associated with enzyme reach, we inspected the “pairwise cross-informing rate” that defines how likely the flux or expression of one reaction is correlated (FDR < 0.05) with the flux or expression of a connected reaction, given the degree of the bridging metabolite that links the two reactions. Indeed, reactions connected by metabolites with a low degree are more likely to cross-inform each other for both flux and expression as well as expression-flux correlation (Fig. 2b).

**Fig. 2:**
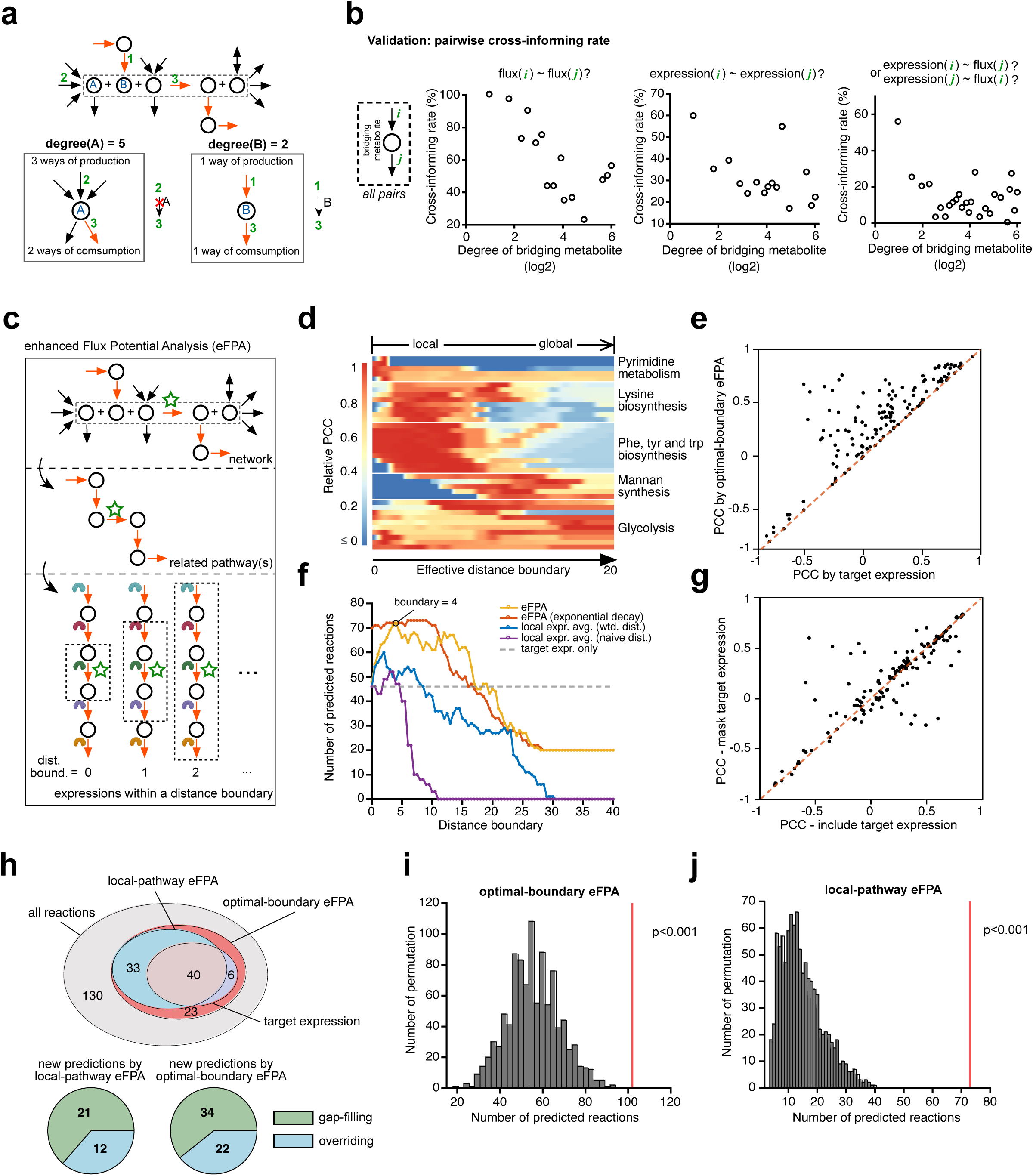
Enhanced flux potential analysis (eFPA) predicts flux by leveraging pathway-level enzyme expression. **a**, Toy network illustrating the degree of bridging metabolites. Circles are metabolites and arrows are reactions. The red arrows indicate a hypothetical pathway for the target reaction inside the dotted box. **b**, Analysis of pairwise cross-informing rate. **c**, Schematic for eFPA. The target reaction is labeled by a green star. **d**, Representative results of eFPA for different reactions. Values are row-normalized flux-rFP PCCs. Rows correspond to different target reactions and columns to the effective distance boundary that indicates the maximum length of pathway(s) integrated (see Materials and Methods). **e**, Comparison of the correlation of flux with target rFP and target expression. **f**, Total number of predicted reactions with different approaches. Local expression average was defined by the average expression of reactions that are connected to target reaction within a distance boundary measured by naïve or weighted distance. **g**, Comparison of local-pathway eFPA predictions with and without inclusion of enzyme levels corresponding to the target reaction. Values are flux-rFP PCCs. **h**, Classification of fluxes predicted by different types of eFPA. **i**, **j**, Randomization test of optimal-boundary eFPA (**i**) and local-pathway eFPA (**j**). The histograms show the distribution of the number of reactions predicted in 1000 permutations and the red line indicates the real observation.

To capture enzyme reach in the yeast metabolic network, we used metabolic network modeling based on flux balance analysis (FBA). Specifically, we adapted our FPA algorithm, which calculates the relative flux potential (rFP) of a target reaction based on enzyme mRNA levels (Yilmaz *et al*., 2020) (Appendix Supplementary Notes). FPA is formulated as an FBA problem whose objective function is the maximal flux of the target reaction and whose maximized objective value is named *flux potential*. The key concept of FPA is to convert relative enzyme levels into weight coefficients such that the flux through a reaction is penalized when cognate enzyme levels are low (Fig. EV2a, Materials and Methods)(Yilmaz *et al*., 2020). FPA uses a distance decay function with which enzyme reach decreases with distance to the target reaction. However, the original FPA did not consider the architecture of the network around a target reaction. Therefore, to better capture enzyme reach, we incorporated bridging metabolite degree to utilize flux routes that are both flux-balanced and low in bridging metabolite degree (Appendix Supplementary Notes) (Fig. EV2b). In addition, we formulated a distance decay function with a *distance boundary* parameter to precisely control the maximum length of integrated pathway(s) (Fig. 2c and Fig. EV2c, Appendix Supplementary Notes). We refer to this algorithm as enhanced FPA (eFPA).

To study enzyme reach in the yeast metabolic network, we performed eFPA using each one of the 232 reactions for which flux was calculated (Hackett *et al*., 2016) as the target reaction. Interestingly, for most reactions, the correlation between rFP and flux varied with distance boundary (Fig. 2d, Fig. EV3). We refer to the boundary where the correlation is maximal as the *optimal distance boundary* and to the corresponding eFPA as “optimal-boundary eFPA”. Overall, the flux of 44% (101/232) reactions could be significantly correlated with rFP (Fig. EV2d). Importantly, optimal-boundary eFPA improved the correlation for most reactions, further confirming that enzyme levels affect metabolic flux of non-cognate reactions (Fig. 2e). The optimal distance boundary of most reactions was between 2 and 6 (Fig. EV3), and the total number of significantly correlated reactions peaked at a distance boundary of 4 (Fig. 2f, yellow curve). As expected, when gene expression was solely considered, *i.e.,* without flux balance or metabolite degree, the correlation with flux diminished (Fig. 2f, blue and purple curve). These observations indicate that enzyme reach occurs mostly locally, within a metabolic pathway. We therefore reasoned that we could generate flux predictions that are less sensitive to the choice of distance boundary, which has to be made a priori, by optimizing the decay function in eFPA. To do this, we used an exponential decay function (Fig. EV2c) and found that this helped to predict relative flux with a broader range of distance boundaries (Fig. 2f, red curve). We refer to the use of exponential decay function and an a priori single distance boundary of 6 as the “local-pathway eFPA”, which is independent of the fitted optimal distance boundaries. Using local-pathway eFPA, we found that only using levels of enzymes catalyzing non-cognate reactions predicted reaction fluxes equally well as when cognate enzyme levels were included (Fig. 2g). Taken together, our results show that metabolic enzymes have a local reach, i.e., their levels affect not only their cognate reactions but also correlate with reactions to which they are coupled by low-degree metabolites and flux balance.

Remarkably, eFPA predicted the relative flux of some reactions only with very large distance boundaries (Fig. 2d and Fig. EV3). For instance, for glutathione synthesis, the levels of enzymes involved in energy generation (glycolysis, oxidative phosphorylation, and the TCA cycle) contributed significantly to accurate flux predictions (Appendix Fig. S2a). We noticed that high (or low) glutathione synthesis usually coincides with high (or low) level of those energy-generating enzymes, which critically contributed to the accurate prediction of flux variations (Appendix Fig. S2b). This observation suggests that glutathione biosynthesis is limited by energy production, which is in agreement with early observations(Yanari *et al*, 1953). Thus, eFPA independently revealed the tight coupling between glycolysis and the synthesis of glutathione solely using enzyme levels.

Overall, eFPA correctly predicted the relative flux of a large proportion of metabolic reactions, including reactions for which enzyme level measurements were not available (*gap-filling predictions*) and reactions for which cognate enzyme levels did not correlate with flux (*overriding predictions*) (Fig. 2h). Finally, a randomization of eFPA by shuffling the reaction labels in the input expression data showed the significance of eFPA predictions, which suggests that enzyme reach is an intrinsic property of the flux-enzyme relationship and not a result of overfitting (Fig. 2i,j).

### Beyond pathway-level coordination

While eFPA predicted relative flux for about half of the reactions, the other half could not be predicted by enzyme levels (Fig. 2h). We hypothesized that some of the fluxes that were not predicted by eFPA could still be indicated from enzyme levels based on flux coupling, where a set of reactions maintain correlated fluxes because the reactions are directly coupled at steady state. An example of such flux coupling is the control of pathway fluxes by a single rate-limiting enzyme, which by itself is not sufficient to predict the flux by eFPA. To empirically derive coupled reactions, we defined the term *flux collinearity* as the absolute PCC between two fluxes in the yeast dataset, where fully coupled reactions (i.e., reactions in a linear pathway) have perfect collinearity with absolute PCC = 1. The flux of many testable reactions (161/232, 69%) was highly colinear with at least one of the 46 correlated reactions (absolute PCC > 0.8, Fig. 3a). Thus, the flux of these non-correlated reactions can in principle be predicted by the levels of at least one enzyme for which the levels correlate with cognate reaction flux, assuming the empirical flux coupling is generalizable in exponentially growing yeast. We refer to these as “indicator enzymes” (Appendix Supplementary Notes, Fig. EV4). As expected, nearly all flux variations predicted by eFPA could also be predicted by an indicator enzyme (96/102, 94%) (Fig. 3a, Venn diagram). Remarkably, 40% of fluxes (65/161) could only be predicted by indicator enzymes, implying that their flux is not controlled by pathway-level coregulation of enzyme expression (Fig. 3a).

**Fig. 3:**
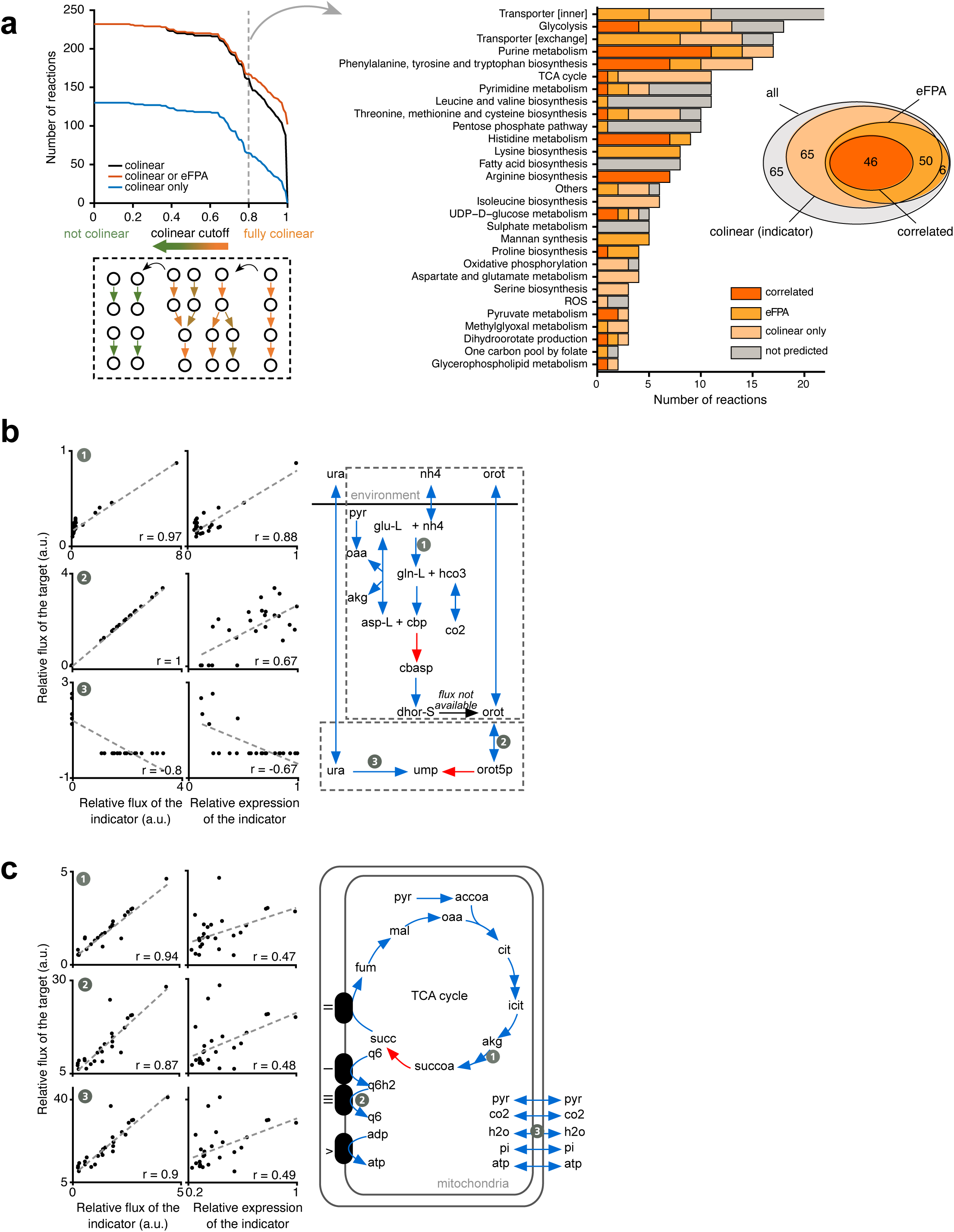
Indicator enzymes complement eFPA and may identify rate-limiting enzymes. **a**, Flux collinearity analysis. Collinearity is defined as the absolute PCC between two reaction fluxes. Given a collinearity cutoff (x-axis), the black line shows the number of reactions that were colinear with any of the 46 correlated reactions. The number of reaction fluxes predicted by either collinearity or eFPA are shown in red, while those only predicted by collinearity are shown in blue. The cartoon illustrates flux coupling between reactions indicated by different levels of collinearity. Pathway annotations of reactions predicted by either correlation, eFPA, or collinearity are shown on the right. **b**, The indicator of pyrimidine metabolism fluxes. In the network cartoon, red arrows refer to the indicator and blue arrows to the target reactions. Two distinct sets of colinear fluxes were identified and are labeled by the two dashed boxes. The correlation between the flux of the target reactions and the flux or expression of the indicator is plotted for representative reactions. **c**, As in (**b**), but for mitochondrial energy metabolism fluxes.

Fluxes in pyrimidine biosynthesis reactions were predicted by two indicator enzymes; flux in the upstream branch that synthesizes orotate correlates with levels of aspartate carbamoyltransferase, while flux in the downstream branch that produces UMP was predicted by orotidine-5’-phosphate decarboxylase levels (Fig. 3b, red arrows). The fluxes of the two branches are not highly colinear because orotate is partially secreted (Hackett *et al*., 2016). Interestingly, the flux of one of the target reactions in which uracil (ura) is converted to UMP (ump) showed a negative correlation with the levels of the indicator enzyme. This may reflect the alternative use of the salvage and *de novo* pyrimidine biosynthesis pathways (Fig. 3b, reaction 3). Importantly, the reactions catalyzed by both indicator enzymes are key rate-limiting steps in UMP synthesis and are known to be regulated both transcriptionally and allosterically in multiple systems(Santoso & Thornburg, 1998; Traut & Jones, 1977). These analyses suggest that yeast pyrimidine biosynthesis is controlled, at least in part, by the levels of two specific rate-limiting enzymes.

Some other indicator enzymes are less likely to catalyze a rate-limiting step. For instance, fluxes in the main energy generating pathways, the TCA cycle and oxidative phosphorylation, were best predicted by levels of succinyl-CoA synthetase, which converts succinyl-CoA to succinate in the TCA cycle, rather than by dehydrogenases such as isocitrate dehydrogenase that catalyze potentially rate-limiting reactions in the cycle (Gabriel *et al*, 1986; Stueland *et al*, 1988)(Fig. 3c). The strong collinearity between TCA cycle flux and other mitochondrial energy production reactions in yeast has been observed previously (Blank & Sauer, 2004). The levels of succinyl-CoA synthetase may serve as a general indicator of the fluxes related to mitochondrial ATP production in yeast.

The flux in reactions of a few pathways, such as the pentose phosphate pathway and fatty acid biosynthesis, could be predicted neither by eFPA nor by indicator enzymes (Fig. 3a). This observation agrees with previous work where the pentose phosphate pathway was found to be regulated allosterically and by substrate concentration (Chubukov *et al*., 2013; Ralser *et al*, 2009; Stanton, 2012). Fluxes in other pathways, such as aspartate and glutamate metabolism and the TCA cycle, could only be predicted by indicator enzymes, which may be due to the prevalence of hub metabolites, the lack of enzyme coregulation in these pathways, or both.

### Enzyme reach in human metabolism

Next, we asked if local-pathway enzyme reach also applies to human metabolism. We performed local-pathway eFPA on a human dataset containing protein and mRNA levels for 32 tissues (Jiang *et al*, 2020) and the human metabolic network model, Human 1 (Robinson *et al*, 2020) (Fig. 4a). Since systems-level flux measurements are not available for human tissues, we could not systematically validate flux predictions like we did with the yeast dataset. However, in the yeast dataset, eFPA captures boundary fluxes, i.e., metabolite exchange with the environment and drainage into biomass, equally well as fluxes within the metabolic network (Fig. 4b, Fig. EV3 and Fig. EV5a). In human tissues, such boundary fluxes represent metabolite secretion or uptake and may provide a proxy for the enrichment of metabolites in different tissues. Therefore, we hypothesized that we could validate boundary flux predictions with available tissue-enriched metabolite annotations (Pang *et al*, 2021).

**Fig. 4:**
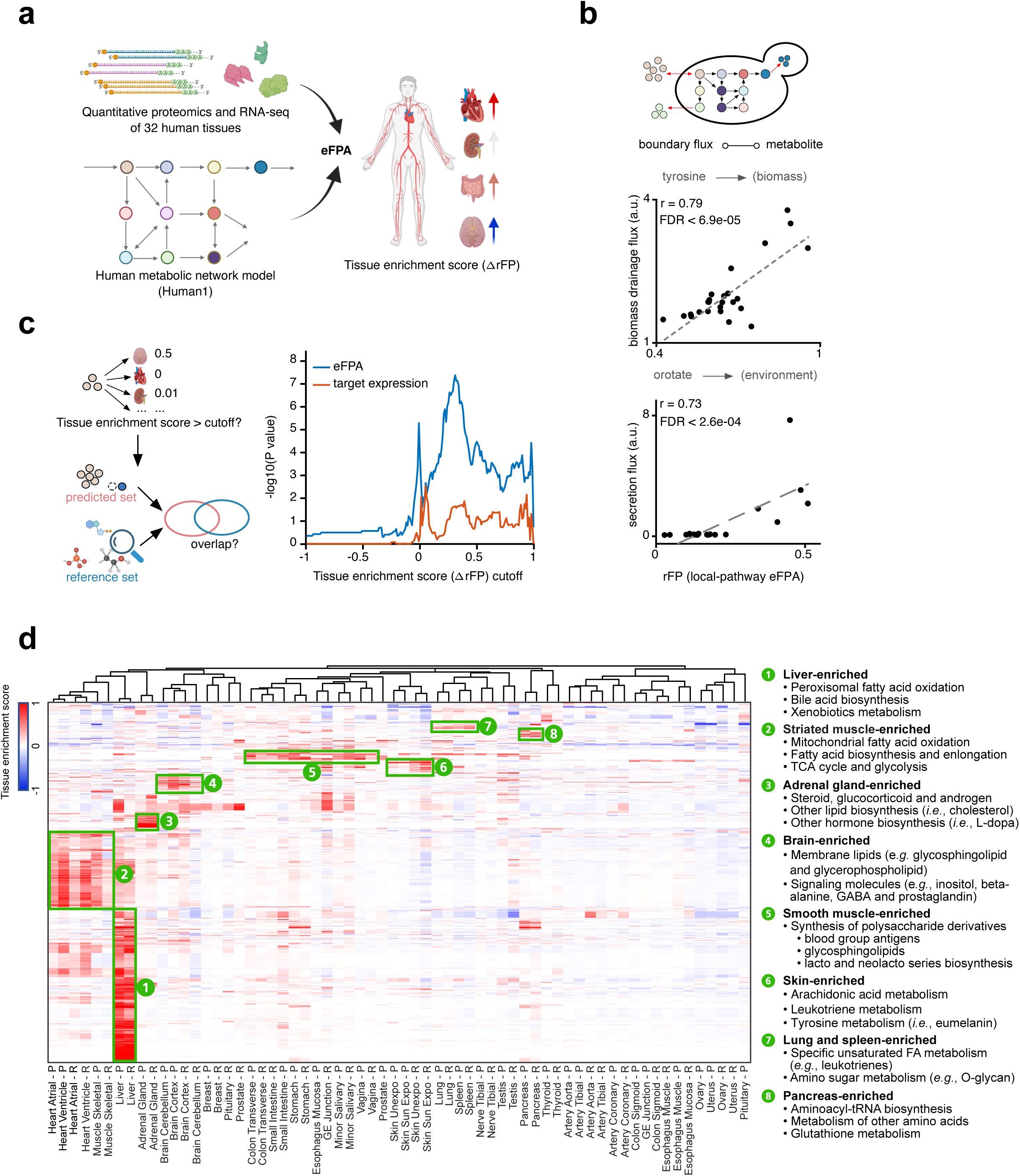
eFPA predictions for human tissue metabolism. **a**, Cartoon of eFPA analysis. rFP values were centered by subtracting the median rFP of a reaction across all 32 tissues, and the difference from median (ΔrFP) was referred to as the tissue enrichment score. **b**, Representative predictions of boundary flux in yeast. The biomass drainage flux of tyrosine and the secretion flux of orotate are shown. **c**, Enrichment p-values (hypergeometric test) between predicted tissue-enriched transportable metabolite set and the reference set under different tissue enrichment score (ΔrFP) cutoffs. **d**, Heatmap of the ΔrFP of 3441 tissue-enriched reactions in 32 human tissues. Tissue-enriched reactions were defined with ΔrFP greater or equal to 0.2 in at least one tissue based on either protein or mRNA expression data as the input. Only high-confidence predictions are shown (see Materials and Methods). Column label shows tissue names with a single letter suffix indicating the prediction was made from protein (-P) or mRNA (-R) data.

To interpret rFPs for tissue enrichment, we defined the tissue enrichment score as the difference of an rFP value from the median rFP of 32 tissues (ΔrFP). For a metabolite, we used the maximum ΔrFP of all relevant transporter reactions that take up or secrete this metabolite. We reasoned that this maximal transport potential can indicate tissue-specific metabolite enrichment. Specifically, a high uptake flux could hint at a tissue’s tendency to assimilate a given metabolite from the bloodstream, leading to its accumulation. On the other hand, a high secretion flux might suggest the tissue actively produces the metabolite, implying a higher concentration of that metabolite in the tissue. These ΔrFP scores were compared to a collection of 827 tissue-enriched metabolite annotations across 17 of the 32 human tissues for which metabolite enrichment annotations are available (Pang *et al*., 2021). We found that local-pathway eFPA better recalled the reference set than using only the levels of relevant transporter(s) (Fig. 4c, Fig. EV5b). This observation suggests that local-pathway enzyme reach also occurs in human metabolism.

So far, our analyses used protein levels as input for eFPA. However, mRNA levels obtained from RNA-seq are much more readily available and, therefore, we repeated the eFPA of human metabolites using mRNA levels(Jiang *et al*., 2020). Protein and mRNA levels performed similarly (Fig. EV5 c, d), and some metabolites were only predicted by mRNA levels, which could be because of the higher coverage of RNA-seq data compared to proteomic data (Fig. EV5 c, e). Therefore, mRNA levels can effectively be used in eFPA.

eFPA generated a comprehensive landscape of human tissue-enriched metabolic fluxes using either protein or mRNA levels as input (Fig. 4d and Appendix Fig. S3, Table EV6). Tissues were frequently predicted to have enriched flux in specific reactions and therefore have specialized metabolic functions (Fig. 4d and Appendix Fig. S3). Many of these are consistent with existing knowledge, such as bile acid biosynthesis in liver (Russell, 2003), active metabolism of membrane lipids and signaling molecules in brain (Barber & Raben, 2019; Hanley *et al*, 1988), and energy metabolism in muscle (Westerblad *et al*, 2010). Our predictions further yielded many hypotheses, for instance, a high metabolic similarity between lung and spleen that was driven by reactions involved in immunometabolism (e.g., synthesis of leukotrienes) (Fig. 4d). Finally, some predictions were strictly derived by local-pathway eFPA. For example, peroxisomal oxidation of palmitoyl-CoA was predicted to be enriched in heart, although this organ was not enriched for the expression of associated enzymes (Appendix Fig. S4). However, this prediction is likely valid because the heart generates energy by burning fatty acids(Lopaschuk & Saddik, 1992). Altogether, these results indicate that eFPA can derive meaningful predictions that are enabled by the inclusion of enzyme reach.

## Discussion

Our findings reveal that metabolic enzyme levels can influence the flux of both cognate and non-cognate reactions. The reach of enzyme levels is usually local, at the level of individual pathways. However, longer distance enzyme reach can also play a role in predicting metabolism. Our eFPA analyses suggest that network architecture plays a key role that can be unveiled by carefully considering the degree of metabolites in the context of flux balance: reactions that form a flux-balanced route via metabolites with a low degree, i.e., that are produced and consumed by few reactions, provide high predictive power for which enzymes can have reach to which reactions. Overall, we uncovered three principles of enzyme reach: a simple correlation where the expression of an enzyme (or transporter) affects flux through its cognate reaction, a distance effect where the coordinated expression of multiple enzymes in a pathway overall affects pathway flux, and another distance effect where the expression of a single indicator enzyme associates with multiple reactions. Our findings highlight that, in addition to flux control by metabolite levels, enzyme expression also plays a key role. Although it is possible that enzyme levels change to accommodate instead of controlling the change of flux, our observation is in agreement with theoretical models of Metabolic Control Analysis (MCA), which proposes that the physiological control of flux is more likely achieved by coordinated changes in multiple enzyme levels and that the controlling effect of each individual enzyme is relatively small (Fell, 2005; Kacser & Burns, 1973).

Importantly, eFPA is based on a reasonably designed mechanistic modeling framework with only one free parameter (distance boundary). This is in contrast to multi-parameter statistical modeling that is prone to overfitting. The unique advantage of eFPA is that it exploits coordinated enzyme expression in the most relevant pathway(s) of a target reaction without prior pathway definitions. eFPA will provide a powerful tool to predict metabolic flux variations for organisms for which both expression profiles and a metabolic network model are available. We demonstrated this with our preliminary application to human tissues, although the absence of flux data prevented the optimization of the distance boundary. We showed that metabolite production and consumption potentials calculated by eFPA statistically relate to experimental metabolite enrichments, the only data type we could use directly, even though these datasets are noisy. Also, the ability of eFPA to infer metabolic flux differences from enzyme expression is clear from the recapitulation of known tissue functions in the form of predicted flux variations.

Finally, the gap-filling predictions by eFPA in yeast (Fig. 2h) indicate that eFPA is robust even in the context of low-coverage data. Therefore, it could be used for interpreting the physiological effects of changes in enzyme expression in various types of omics data, including single-cell RNA-seq and proteomics. Ultimately, this would facilitate a deeper understanding of metabolism in diverse conditions and at the level of individual cells.

## Materials and Methods

Brief descriptions of the main methods used in this study are included here. Details for each of the following sections are provided in Appendix Supplementary Methods, following the same order of related sections listed here as well as in the main text.

### Processing of SIMMER dataset

Fluxomic and proteomic data from Hackett *et al*.(Hackett *et al*., 2016) were transformed to facilitate correlation analysis and metabolic network modeling. We adjusted raw (absolute) flux values by dividing them by the corresponding growth rate to obtain the flux levels used throughout this study. To be consistent with the Flux Potential Analysis (FPA)(Yilmaz *et al*., 2020), where unscaled expression levels are used, we exponentially transformed the log2-scale proteomics data to obtain the unscaled relative abundances (referred to as *protein level*). Finally, to correlate the flux of a reaction with expression level of enzymes associated with it, the *protein levels* need to be converted to a single value that represents the expression level of the reaction according to the Gene-Protein-Reaction (GPR) associations(Yilmaz *et al*., 2020). We followed a method described previously(Yilmaz *et al*., 2020) that produces a normalized, reaction-level, relative expression across conditions that varies from 0 to 1. We refer to this value as *relative expression of reaction*.

The flux and protein data were mapped to the most recent consensus metabolic model of yeast (yeastGEM_v8.3.5(Lu *et al*, 2019)). Out of the 233 flux values, 232 were mapped to the corresponding reactions, with one reaction, r_1099, discarded because of changes in the reaction formula in the new yeast model. Proteins encoded by 486 genes in the model were found to be quantified by proteomics, accounting for 42% of all model genes and associated with 809 reactions (20%). Out of these 809 reactions, 658 had complete expression measurement (i.e., all associated proteins were quantified). A total of 156 reactions had both the flux and enzyme expression levels determined. These reactions were used in the correlation analysis. Such dual-omics dataset with fluxes and enzyme expression levels was generated for each one of the 25 conditions where media composition and dilution rate were varied.

### Correlation analysis of flux and enzyme expression level

For each of the 156 reactions whose flux and expression data are both available, we calculated the Pearson correlation coefficient (*PCC*) between *relative expression of reaction* and *flux* using the 25 conditions as 25 datapoints. The *p-value* of each correlation was calculated based on a two-tailed hypothesis test using the *corr* function in MATLAB 2019a and adjusted for multiple testing by *mafdr* function of MATLAB with the ‘*BHFDR’* method (Benjamini and Hochberg (BH) FDR correction). The resulting FDR values were used to evaluate the correlation. A correlation was considered significant if the FDR is less than or equal to 0.05.

### Analysis of pathway-level coexpression

We derived the *PCCs* of *relative expression of reaction* (over the 25 conditions) for every pairwise combination of reactions in a defined pathway (Fig. EV1e and Table EV2) and took the median value of these *PCCs* to define the strength of pathway coexpression. Similarly, we defined the strength of pathway-level flux-expression correlation as the median of the flux-expression *PCCs* for all reactions in a pathway. Pathway-level coexpression patterns were defined as the median of *relative expression of reaction* over pathway reactions which formed a 25-element vector for each studied pathway.

### Analysis of pairwise cross-informing rate

We considered all connected reaction pairs with given flux and/or protein levels for the cross-informing rate analysis. For instance, to calculate the flux-flux cross-informing rate, we collected every connected reaction pair with fluxes determined for both reactions. We calculated *PCC* and *p-values* for each collected pair and considered reaction pairs with FDR less than 0.05 and *PCC* greater than 0 as *cross-informed*. The cross-informing rate was defined as the proportion of cross-informed pairs in a given set of pairs. The sets of pairs were determined based on the network degrees of bridging metabolites, i.e., metabolites that connect the paired reactions. Importantly, due to the limited number of data points (i.e., only 232 reactions with estimated fluxes), we grouped the pairs with approximate bridging metabolite degrees in the calculation of cross-informing rate. Each group produced a single data point in Fig. 2b in which the x-axis refers to the average bridging metabolite degree of pairs in the group and y-axis refers to the proportion of *cross-informed* reaction pairs, i.e., the cross-informing rate for the group.

### Enhanced Flux Potential Analysis (eFPA)

FPA (and eFPA) is a specific flux balance analysis (FBA) problem that calculates the maximum flux potential (FP) of a target reaction under certain constraints that address relative expression of reactions and their distance from the target(Yilmaz *et al*., 2020). The mathematical details of FPA can be found in Appendix Supplementary Methods and in our previous publication(Yilmaz *et al*., 2020). In brief, as in a regular FBA problem, FPA is done under the steady-state assumption with reaction fluxes constrained between prescribed upper and lower boundaries. Specifically, a weighted sum of flux in the network is further constrained to be less than or equal to a constant that is referred to as *flux allowance* while the weight of each flux is inversely proportional (but not linearly, see the next section) to the *relative expression of (the corresponding) reaction.* The flux of the target reaction is selected as the objective function, which is maximized to find FP as the objective value. Since the flux allowance is a constant, reactions with smaller weight coefficients (i.e., higher relative expression) in the weighted sum are more likely to carry larger flux to maximize the flux left for the target reaction, thereby conveying the influence of their expression changes to the flux potential of the target. The enhanced flux potential analysis (eFPA) algorithm stays the same as original FPA except for the new distance decay functions and the use of weighted metabolic distance instead of naïve metabolic distance (see below).

### Weight coefficients and distance decay function

The key component of FPA is the calculation of the *weight coefficients*. Weight coefficient is proportional to the reciprocal of the *relative expression of reaction* and is scaled by a distance decay function. The distance decay function represents the decrease of the influence of network reactions (enzyme reach) on FP as their distance from the target reaction increases. Thus, the distance function downscales the weight coefficients as the distance to the target increases, which would allow a reaction to take large flux values without significantly affecting target FP calculations even if its expression coefficient is high (i.e., if it is poorly expressed). The eFPA algorithm employs two redesigned distance decay functions, distinct from that in the original FPA. Please refer to Appendix Supplementary Methods for mathematical details.

To integrate the bridging metabolite degree (an important indicator of the network architecture) (Fig. 2a) with eFPA, we weighted the metabolic distance between two adjacent reactions based on the number of connections to the metabolites that connect them (Fig. EV2b), such that the distance between reactions connected by hub metabolites is upscaled. Thus, the distance between a pair of reactions can be greater than the original distance that is the number of reactions between them plus one. We refer to the new distance measure as *weighted metabolic distance*. Weighted metabolic distance effectively encourages the reach of influence from enzymes in linear pathways to the target reaction (i.e., connected by metabolites of lower network degrees), thus achieving the integration of enzyme changes of interest. The distance boundary parameter in eFPA is measured in the scale of weighted metabolic distance. To relate this parameter to the actual length of the integrated pathway for interpretation, we further converted it to an interpretable metabolic distance from the target (i.e., the maximum distance of integrated reactions to the target) that was used for data visualization in Fig. 2d and Fig. EV3. Please refer to Appendix Supplementary Methods (sections *Mathematical formulation of Flux Potential Analysis (FPA)*, *Weight coefficients and distance decay function*, *Weighted metabolic distance* and *Calculation of effective distance boundary*) for the mathematical details.

### eFPA of SIMMER dataset

As in our previous FPA analysis(Yilmaz *et al*., 2020), eFPA does not rely on any quantitative boundary constraints as we are focused on the effect of enzyme expression changes. All available nutrients based on the definition of culturing media were made freely exchangeable by setting the lower boundary of pertaining exchange reactions to –1000. These nutrients include phosphate, glucose, ammonium, uracil and leucine (Table EV5). No non-growth-associated maintenance (NGAM) was imposed (lower boundary to 0) during eFPA. This constrained model was used in all eFPA analyses.

To qualitatively account for media differences across conditions, we blocked the uptake of unavailable nutrients in each condition. For example, uracil and leucine uptakes were blocked in the eFPA of phosphate-limiting, carbon-limiting and nitrogen-limiting conditions. In addition, to address the extreme low abundance of nutrients (e.g., glucose concentration is over 20-fold lower in carbon-limiting conditions), we set an arbitrarily large gene expression coefficient on the pertaining exchange reactions when applicable. Please refer to Appendix Supplementary Methods for details.

eFPA was performed by a modified version of the generic eFPA function (https://github.com/WalhoutLab/eFPA) to enable highly parallel computation in a computer cluster. The distance boundary parameter was changed according to the question of interest as indicated in the text. All 232 reactions (Table EV2) with determined fluxes in SIMMER dataset were analyzed with eFPA.

Other details regarding parameterization, analysis and interpretation of eFPA in yeast can be found in Appendix Supplementary Methods (sections *Correlation analysis of eFPA results and flux data*, *Titration of the distance boundary*, *Calculation of effective distance boundary*, *Deciphering prediction mechanisms of eFPA* and *eFPA on the drainage flux of biomass precursors*).

### Randomization test of eFPA

To assess the statistical significance of eFPA modeling, we shuffled the rows (reaction labels) of the *relative expression of reaction* matrix (658 reactions by 25 conditions), i.e., randomized the association between the expression levels and reaction labels. After shuffling, eFPA was performed with randomized data following the same procedure as described above. This randomization was performed for 1000 times. It is noteworthy that we were unable to perform a greater scale of randomization due to the overwhelming computational demand.

### Processing of human tissue dataset

The quantitative proteomics and transcriptomics of human tissues contain RNA and protein levels of more than 12,000 genes across 32 normal human tissues quantified based on 201 individual primary samples(Jiang *et al*., 2020). We analyzed all 32 tissues using the *tissue median* provided in the referred study. We rescaled both RNA and protein tissue medians to make them suitable for system-level modeling, including the conversion of log2-scale data to unscaled values and a variance-stabilizing transformation based on Tissue Specificity score reported in the referred study(Jiang *et al*., 2020). Please refer to Appendix Supplementary Methods, section *Processing of tissue expression data*, for details about the data transformation.

### Processing of tissue-enriched metabolite set

The tissue enriched metabolite sets were obtained from MetaboAnalystR software (version 4.93, www.metaboanalyst.ca)(Pang *et al*., 2021). There were 73 tissue-enriched or subcellular-enriched metabolite sets in the original data table. Tissue labels in MetaboAnalystR metabolite set were manually matched with the 32 tissues measured in Jiang *et al*. (Jiang *et al*., 2020) (Table EV7). In the end, 331 metabolites from the human metabolic model were defined with an enrichment in at least one of 17 matched tissue types, resulting in 827 tissue-metabolite pairs in total (Table EV8).

### Human metabolic network model

The newest consensus human metabolic model, Human1 (version 1.5.0), was downloaded from metabolicatlas.org(Robinson *et al*., 2020). To increase the numerical stability, the stoichiometry matrix of the model was adjusted to avoid reactions with overly large coefficients or small flux capacities. To model the differential nutrient availability in blood stream, a set of specialized uptake reactions were added to control the mass influx of each type of imported nutrient using a flux balance method that we developed previously(Yilmaz *et al*., 2020). Detailed descriptions on model modifications can be found at Appendix Supplementary Methods, section *Human metabolic network model*.

### eFPA analysis for human tissues

To comprehensively model the tissue metabolism in humans we followed the modeling pipeline MERGE, which we previously developed for modeling *C. elegans* tissue metabolism(Yilmaz *et al*., 2020). In brief, two stages of modeling were performed in a sequential manner. In the first step, a semi-quantitative modeling of on/off status and direction of reaction fluxes was performed using iMAT++ algorithm(Yilmaz *et al*., 2020). This step is to globally fit the distinct expression levels of enzymes in tissues to the metabolic network, to derive (1) a flux distribution that we named as Optimized Flux Distribution (OFD) and that can be used to assign the flux directions for reversible reactions; and (2) a tissue-specific metabolic network in which inactive reactions that cannot carry flux according to flux variability analysis(Yilmaz *et al*., 2020) were removed from the network. After iMAT++ modeling, the eFPA analysis was performed on the derived tissue networks. In addition, the rFP predictions generated from eFPA was overlayed with OFD to derive high-confidence predictions with reaction directionality. This step is important for the interpretation of eFPA predictions of reversible reactions that show high flux potential in both directions, such that the direction predicted by iMAT++ is chosen as the most likely metabolic function with high potential. Together, this tissue modeling pipeline provides a comprehensive collection of high-quality predictions about tissue-enriched metabolic fluxes.

Two sets of tissue networks were built for the 32 tissues based on either proteomics data (protein-based tissue network; PTN) or RNA-seq data (RNA-based tissue network; RTN). PTN and RTN were used in eFPA based on proteomics and RNA-seq data, respectively. The decay function of eFPA algorithm used in human tissue modeling was the exponential decay with a distance boundary of 6 (i.e., the local-pathway eFPA was used), and the distance measurement is the weighted metabolic distance. When not specified (i.e., referred to as *“eFPA”* or *“Protein eFPA”*), the input data for eFPA were protein levels of genes that are commonly detected by both proteomics and RNA-seq (12,121 genes, covering 2869/3627, 79%, genes in the Human 1 model). For eFPA using RNA expression as input (referred to as *“RNA eFPA”* in figures and texts), we performed eFPA using either commonly detected genes (12,121, labeled as *“common genes”* or not specified) or all detected genes by RNA-seq (19,273, labeled as *“all genes”*).

To derive the tissue-metabolic landscape, we analyzed all regular reactions in human 1 model except for transport, exchange, and custom uptake reactions (see above). To model the tissue-enriched metabolites, we performed eFPA on transporter reactions that take up or secrete metabolites from or into extracellular space, respectively. The computation was performed in Massachusetts Green High Performance Computing Center (GHPCC) with 512 cores. To exemplify the computational performance, eFPA of all internal regular reactions (7,248) and 32 tissues took less than 30 minutes.

Detailed parameters, settings and modifications can be found in Appendix Supplementary Methods, sections from *Building human tissue networks* to *eFPA analysis: boundary reactions*.

### Linking boundary flux predictions to tissue-enriched metabolites

eFPA produced an rFP matrix with columns representing the 32 tissues and rows the reactions (i.e., internal reactions and transporters). We next row-wise centered this matrix by subtracting the row median, so that we obtained the delta rFP from the median (ΔrFP). The maximum ΔrFP among the transporter reactions that import or export a target metabolite was defined as the tissue enrichment score of the metabolite. This yielded predictions for 289/331 metabolites that we had reference data for. These 289 metabolites accounted for 740/827 metabolite-tissue pairs that were annotated as tissue-enriched, and were used in further evaluations. To convert the ΔrFPs into binary calls of tissue enrichment (*enriched* or *not enriched*), we set a threshold on the ΔrFPs, where metabolite-tissue pairs with ΔrFPs higher than this cutoff were labeled as *tissue enriched*. Hypergeometric test was used to check if the predicted *tissue enriched* metabolite-tissue pairs significantly overlap with the reference *tissue enriched* metabolite-tissue pairs under various ΔrFP threshold. To compare the ΔrFP values for known tissue-enriched metabolites with that for all metabolites (approximately null distribution for non-tissue-enriched metabolites), the ΔrFP values for all transportable metabolites (i.e., the maximum ΔrFP of all pertaining transporter reactions given a metabolite of interest) in the network (n=1,379) were computed. We compared this overall distribution with the ΔrFP distribution of the 740 reference tissue-metabolite pairs using boxplots. We also tested whether the median ΔrFP of tissue-enriched metabolites is greater than the median of the background via rank-sum test.

### Clustering rFPs of internal reactions

To generate Fig. 4d, reactions were filtered unless their ΔrFP were greater than 0.2 (tissue-enriched) or lower than –0.2 (tissue-depleted) in at least one of the 32 tissues. To address the cases when eFPA predicts tissue enrichment for both directions of a reversible reaction, we selected the eFPA predictions based on the relevant flux distribution (OFD) predicted by iMAT++ (see above). The reactions that passed these criteria were merged at the end. We termed these final filtered predictions *high-confidence predictions of human tissue metabolism* and generated the clustergram with them using the *clustergram* function in MATLAB with ‘cosine’ distance for both row and column clustering. These high-confident predictions were starred (i.e., labeled with “*”) in their reaction ID in Table EV6.

### Clustering tissue-enrichment of subsystems

To visualize the subsystem-level metabolic specialization in tissues, we generated a matrix where rows are subsystems and columns are the 32 tissues. The values in the matrix are number of tissue-enriched reactions (ΔrFP greater than 0.2) assigned to each subsystem and tissue. The matrix was row-wise normalized by dividing all values with the maximum, which yielded the relative tissue-enrichment for subsystems shown in the heatmap (Appendix Fig. S3).

## Data availability

Core data are available in the main text or the Appendix. The complete datasets and codes to reproduce the presented work can be found at https://github.com/WalhoutLab/eFPA. A tutorial for applying eFPA is also included in the Github repository.

## Acknowledgments

We thank members of the Walhout lab for supports and discussions on this project. We thank Caryn Navarro, Hefei Zhang, Job Dekker, Olga Ponomarova, Robert Brewster and Shivani Nanda for discussions on the manuscript. This work was supported by a grant from the National Institutes of Health GM122502 to A.J.M.W.

## Author contributions

X.L., L.S.Y. and A.J.M.W. conceived the project and wrote the manuscript. X.L. and L.S.Y. developed the eFPA algorithm. X.L. performed all investigations under supervision of L.S.Y. and A.J.M.W.

## Supporting information

Appendix

Supplementary Methods

Supplementary Notes

Supplementary Figures 1-4

## Expanded view figures

Figures EV1-4

## Tables

Tables EV1-9

## Expanded view figure legends

**Fig. EV1:**
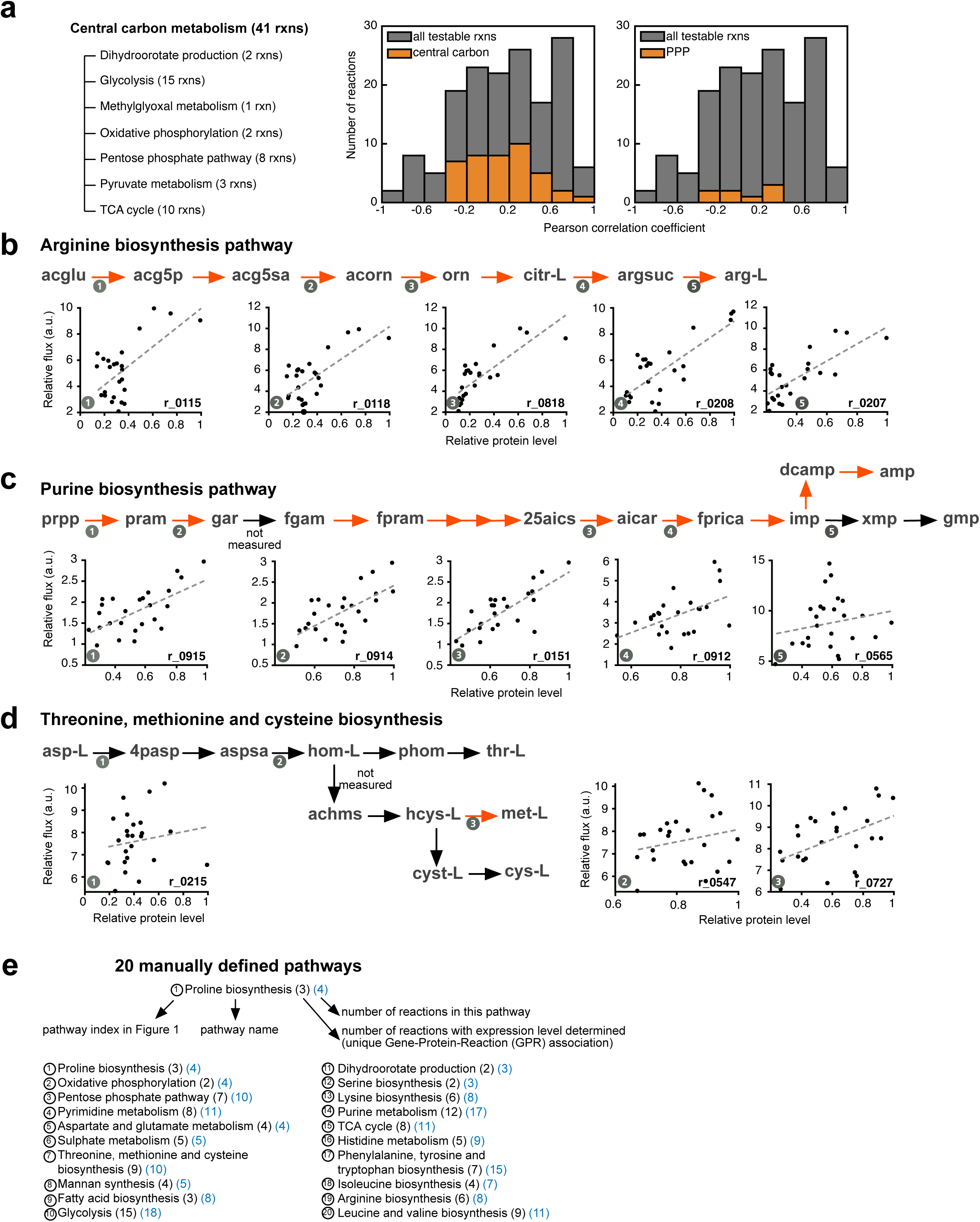
Analysis of flux-enzyme level correlation in metabolic pathways. **a**, The *PCC* distribution for reactions of central carbon metabolism. The distribution of reactions of the pentose phosphate pathway (PPP) is shown on the right. **b**,**c**,**d**, overlay of the flux-enzyme level correlations on selected metabolic pathways. The three selected pathways include ones that are fully (**b**) or partially (**c**) composed of correlated reactions, and one that has only one correlated reaction (**d**). Orange arrows indicate reactions that show significant correlation (FDR < 0.05, *PCC* > 0) and black arrows indicate uncorrelated reactions. **e**, Twenty manually-defined pathways that are labeled by their indices in Fig. 1e,f.

**Fig. EV2:**
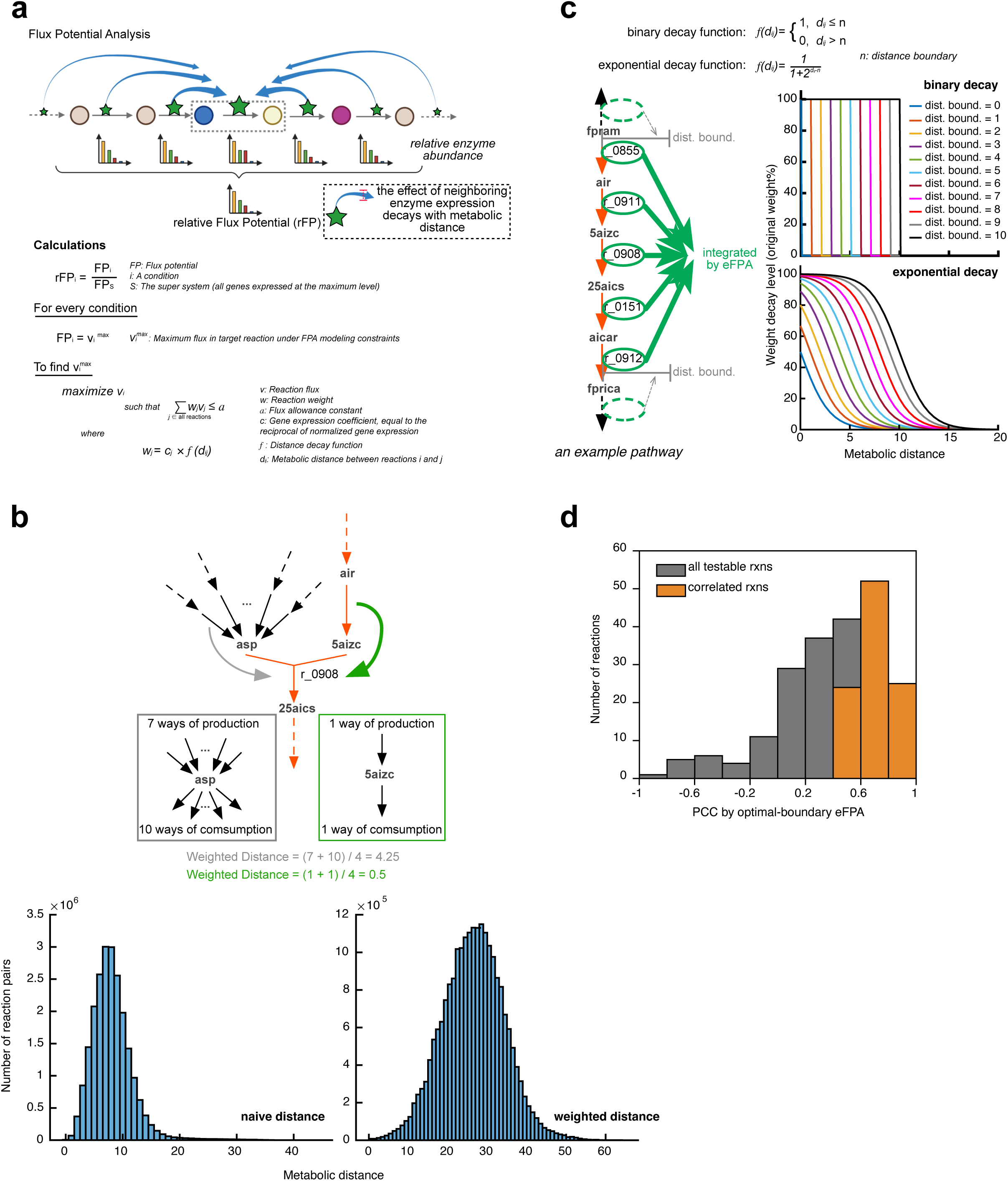
Principles of FPA and eFPA. **a**, Schematic of Flux Potential Analysis (FPA). FPA and eFPA integrate the levels of enzymes that catalyze reactions surrounding a target reaction to predict relative flux potential (rFP) of the target reaction. The contribution of surrounding enzymes can be tuned by a distance decay function such that FPA is versatile to integrate expression information from a local subnetwork to the entire network. The mathematical formulation of FPA is briefly summarized (also see Appendix Supplementary Notes). **b**, The weighted metabolic distance. A cartoon illustrating the calculation of weighted metabolic distance is shown on the top. On the bottom, the distributions of naïve (unweighted) and weighted metabolic distance are shown for all reaction pairs in the yeast metabolic model. **c**, The distance decay functions of eFPA. The formulas of binary (used in optimal-boundary eFPA) and exponential decay functions (used in local-pathway eFPA) are shown in the figure. The left panel shows a cartoon to illustrate the concept of distance boundary. The right panel shows the decay curve of the two functions. **d**, The distribution of *PCC* between experimental flux and rFP predicted by optimal-boundary eFPA for the 232 reactions with flux estimates. Significantly correlated reactions (FDR < 0.05, *PCC* > 0, min-max rFP range > 0.2) are colored in orange.

**Fig. EV3:**
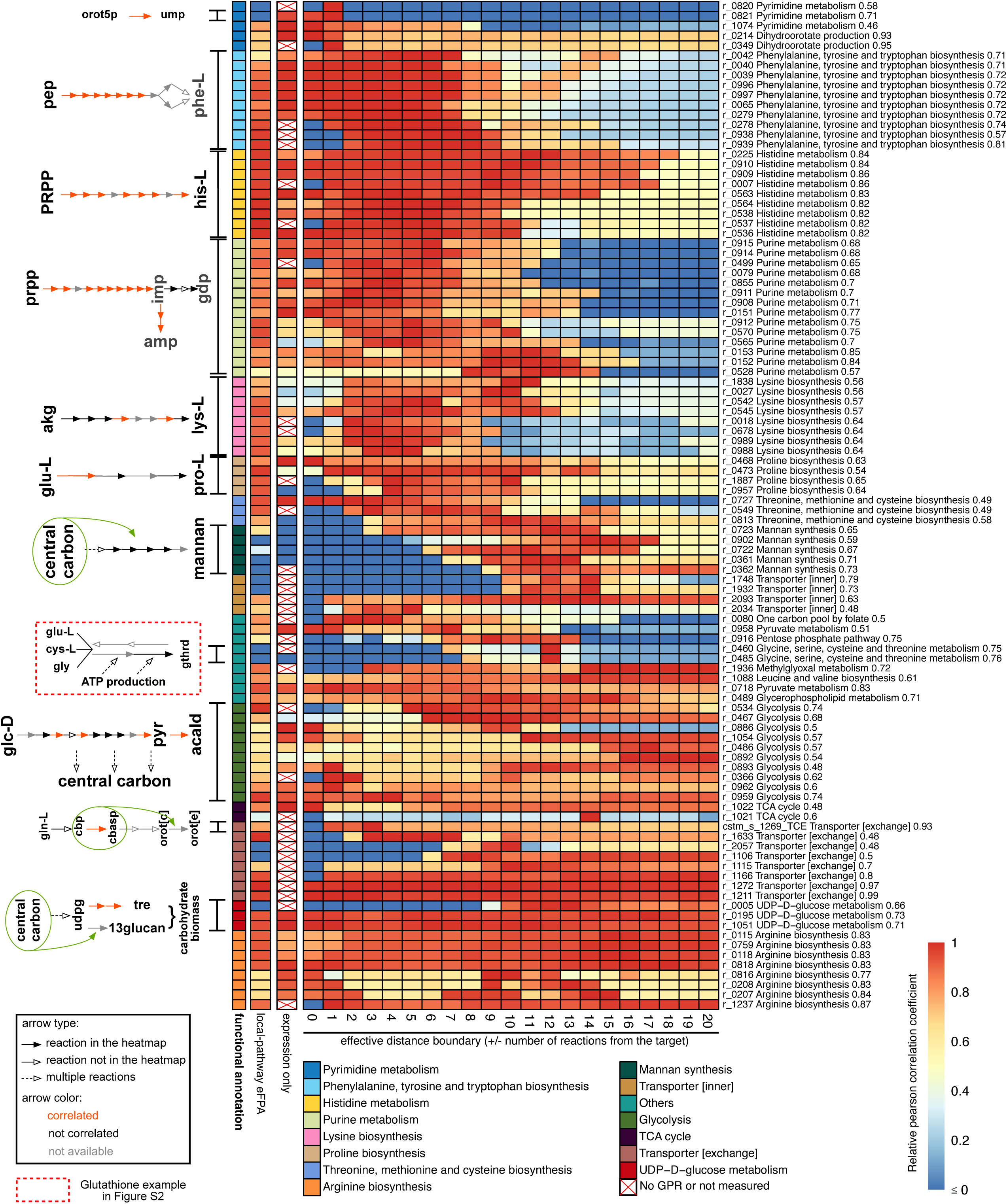
Heatmap of PCC between flux and rFP with a titration of distance boundary. The relative PCC (row-wise normalized by dividing each value by the row maximum) for 102 predicted reactions. The numbers on x-axis indicate the effective distance boundary, which represents the calculated actual length of the integrated pathway based on the distance boundary parameter in the scale of weighted metabolic distance. Target reactions (rows) are arranged based on their position in the pathway they are associated with, and reaction IDs are indicated on y-axis together with the associated pathway and the optimal-boundary PCC. Significant contributors (e.g., relevant local pathways) to eFPA analysis of some targets are depicted on the left.

**Fig. EV4:**
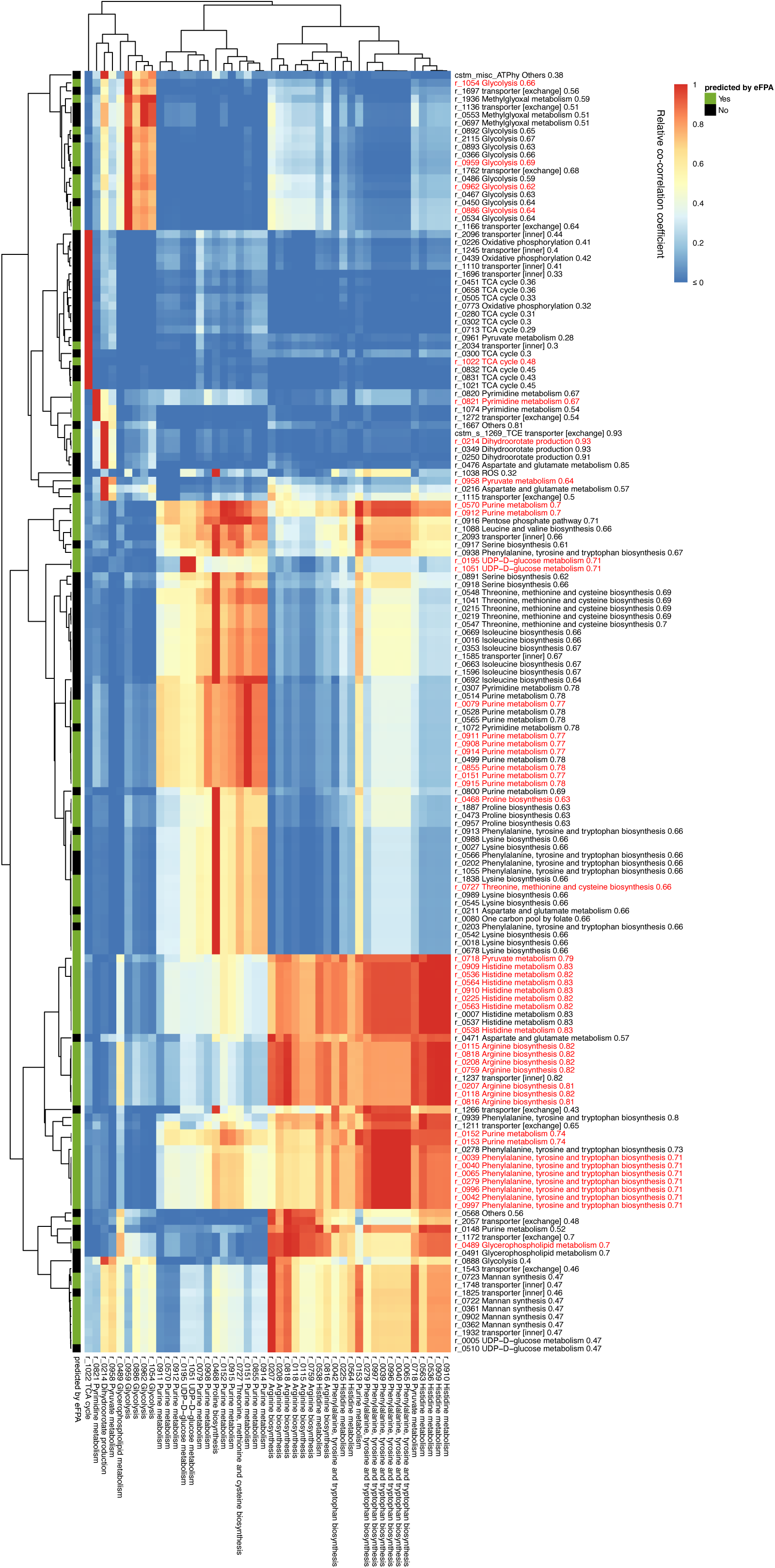
Heatmap of co-correlation coefficients to identify reactions predicted by indicator enzymes. Reactions (targets, rows) whose collinearity with any of the 46 correlated reactions (indicators, columns) is greater than 0.8 are shown. The co-correlation coefficient is defined as the product of the PCC between indicator and target fluxes and the PCC between target flux and indicator enzyme expression (see Appendix Supplementary Notes). This metric indicates how well the enzyme levels of an indicator reaction can be used to predict the target flux variations. The coefficients were row-wise normalized (i.e., by dividing the values with the row maximum) for visualization. The maximum co-correlation coefficient of each row is indicated next to the reaction name. Target reactions that belong to the set of 46 correlated reactions are highlighted in red on the y-axis.

**Fig. EV5:**
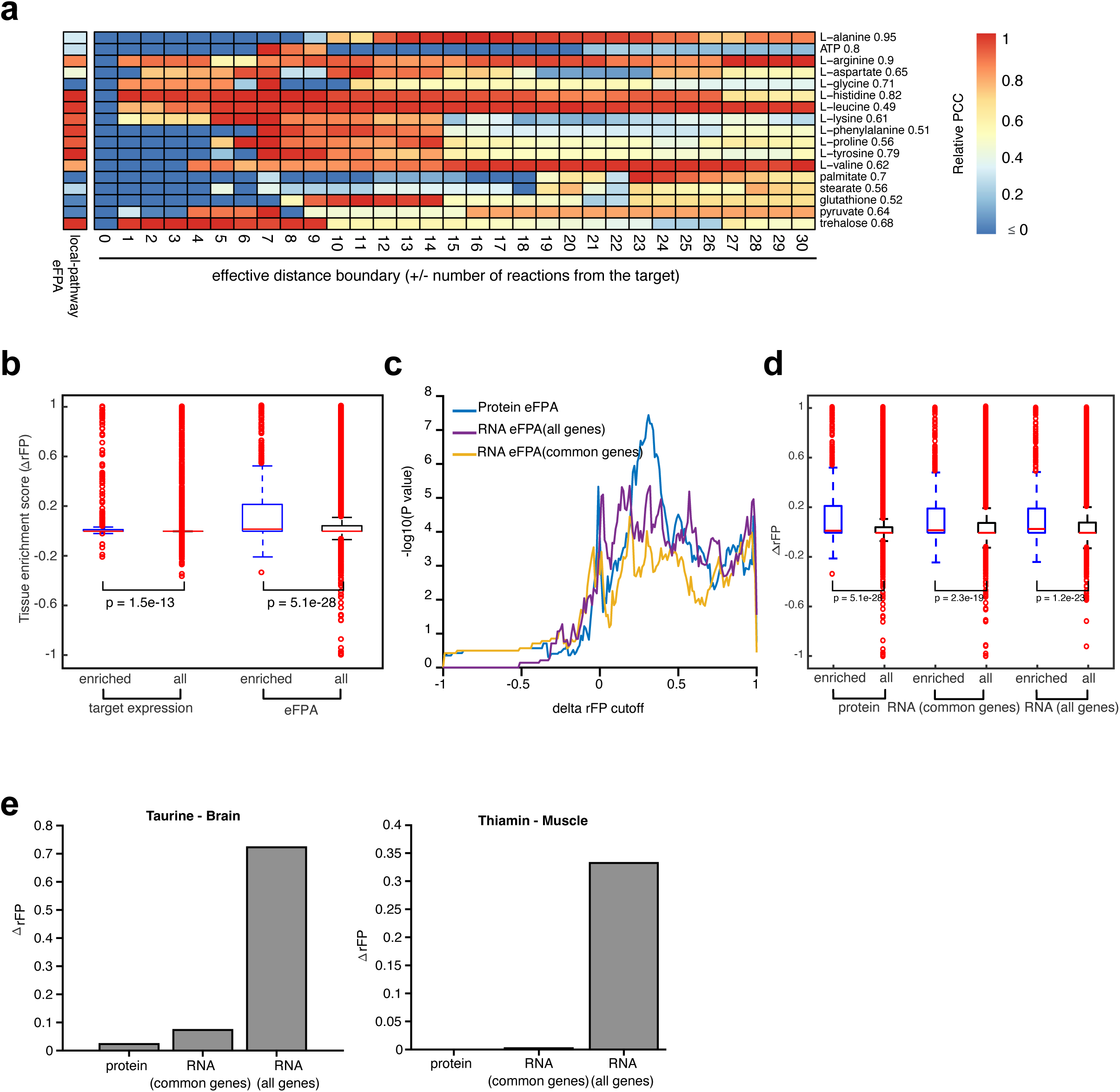
Investigating local-pathway enzyme reach by predicting boundary fluxes. **a**, Heatmap of flux-rFP PCC with a titration of distance boundary for the drainage fluxes of biomass precursors in the yeast model, which serve as boundary fluxes for these metabolites(Hackett *et al.*, 2016). The relative (row-wise normalized) PCCs of 17 predicted (FDR < 0.05, PCC > 0, min-max rFP range > 0.2) reactions are shown in the heatmap. The layout of the heatmap is identical to Fig. EV3. **b**, Boxplots of metabolite tissue enrichment scores (ΔrFP) based on transporter flux potentials calculated by transporter expression only or by eFPA. The scores for the reference tissue-enriched metabolites (*enriched*) were compared with that of all transportable metabolites in Human 1 (*all*). P-values indicate the significance of difference between the median score of enriched metabolites and that of all metabolites (rank-sum test). Both the box statistics and p-values show a larger increase of *enriched* with respect to *all* in the case of eFPA. **c**,**d**, Comparison of eFPA results obtained using protein and mRNA levels. Common genes and all genes refer to the gene set that is shared in protein and RNA-seq data and the set of all genes covered in the RNA-seq data alone, respectively. Results were evaluated both by the overlap with the reference set (**c**) and by the levels of tissue-enrichment scores (**d**). **e**, Two examples showing that the higher coverage of RNA-seq data enabled useful predictions that were missed in the protein data modeling.

