## Appendix for "Linking enzyme expression to metabolic flux"

**This PDF file includes:**

Supplementary Methods
Supplementary Notes
Supplementary Figures 1 to 4

**Other Supplementary Information for this manuscript include the following:**

Table EV 1 to 9

### Supplementary Methods

Analysis on SIMMER Dataset

*Processing of expression and flux data*

Fluxomic and proteomic data was obtained from Hackett *et al*. [1]. The dual-omics data were measured for 25 conditions where media composition and dilution rate were varied. The dataset provides quantitative proteomics for a total of 1187 proteins, in the form of log2 relative abundance with respect to a common reference (an internal $^{15}$N-labeled reference sample). To be consistent with the Flux Potential Analysis (FPA) [2], where unscaled expression levels are used, we exponentially transformed these values to obtain the unscaled protein abundances (referred to as ***protein level***) (Equation S1).

$$protein\ level = 2^{reported\ value} \quad \text{S1}$$

Fluxomic part of the reference dataset provides the upper and lower limits of metabolic fluxes as determined by Flux Variability Analysis (FVA) in a yeast genome-scale metabolic network model (MNM) [3] which was constrained with experimentally determined fluxes including the nutrient uptake rates, by-product secretion rates, and biomass precursor production rates. The flux of 233 reactions was considered as 'determined' because they were well constrained based on FVA ((FVA range) / (midpoint of FVA range) < 0.3). We used the midpoint of these 233 reactions as the input flux value in our downstream analyses.

It is worth noting that Hackett *et al.* also reported an optimal flux value for each of the 233 reactions by minimizing the global difference between fitted fluxes and experimental measurements in a quadratic programming (QP) problem. We found that the QP-flux showed similar results with the midpoint of FVA interval, so we used the midpoints in our final analysis to avoid potential complications introduced by QP fitting.

It is critical to properly normalize the flux values, as we should compare protein levels and flux levels within the same quantitative category. For instance, the input flux measurements in the reference study are in the unit of *moles/hr/mL cells*. In contrast, the input protein levels are unit-free and represent the normalized amount of a given protein that was determined relative to the total protein content (i.e., equal amounts of labeled and unlabeled total protein were mixed in the preparation of proteomics samples). Therefore, a comparable flux quantity would be the flux level normalized to total flux or some internal control flux. For yeast cultured in chemostats, metabolic fluxes scale with the dilution rate (equal to growth rate) that determines the input and output rate in the final steady state. We therefore reasoned that flux levels can be normalized by the growth rate to obtain adjusted fluxes that are comparable to the protein data. Thus, we divided the input flux by the corresponding growth rate ($h^{-1}$). Since the growth rate in Hackett *et al*. was determined by an optical density measurement that is proportional to total cell volume instead of total cell weight and, consistently, the unit of the flux values is *moles/hr/mL cells*, we normalized the input flux by the growth rate without unit conversion (Equation S2). Hereafter this adjusted flux level is simply referred as *flux*. We will focus on comparing the relative levels of flux across conditions with enzyme expression.

$$adjusted\ flux = \frac{flux}{growth\ rate} \quad S2$$

The flux and protein measurements were mapped to the most recent consensus metabolic model of yeast (yeastGEM_v8.3.5 [4]). Out of the 233 flux measurements, 232 were mapped to the corresponding reactions, with one reaction, r_1099, discarded because of changes in the reaction formula in the new yeast MNM. Proteins encoded by 486 genes in the model were found to be quantified by proteomics, accounting for 42% of all model genes and associated with 809 reactions (20%). Out of these 809 reactions, 658 had complete expression measurement (i.e., all associated proteins were quantified). A total of 156 reactions had both the flux and enzyme expression levels determined. These reactions were used in the correlation analysis. Such dual-omics dataset with fluxes and enzyme expression levels was generated for each one of the 25 conditions where media composition and dilution rate were varied.

*Correlation analysis of flux and enzyme expression level*

To correlate the flux of a reaction with expression level of enzymes associated with it, the enzyme levels need to be converted to a single value that represents the expression level of the reaction. While this is straightforward for reactions associated with single enzymes encoded by single genes (i.e., reaction expression level = enzyme expression level), in the case of reactions associated with multiple enzymes or enzyme complexes encoded by multiple genes, enzyme abundances need to be processed using Gene-Protein-Reaction (GPR) associations to form an overall expression level of a reaction. We calculated the expression level of reactions from GPR rules following the methods in our previous work [2] using the protein levels mentioned above as the input. In brief, for protein complexes, we used the minimal expression level of all subunits to represent the expression of the complex; while in the case of isoenzymes, we used the total abundance to represent the overall expression. In the end, the expression level of each reaction in each condition was normalized by the maximum expression level across the 25 conditions to obtain the ***relative expression of reaction***, a quantity that varies from 0 to 1. To avoid potential complications arising from detection issues, we discarded reactions for which expression levels were not available for all of the associated enzymes.

It is noteworthy that, although the proteomics data is presented as a relative quantity (fold change), we added up the levels of isoenzymes to be consistent with the previous rules [2]. This may cause some distortion of the relative expression levels. For example, in a reaction associated with two isozymes, if one of the isozymes is highly expressed and increases 2-fold from one condition to another while the other isozyme is lowly expressed and is unchanged between these conditions, the expected change in the *relative expression of reaction* should reflect the 2-fold increase of the highly expressed isozyme since we use the summation rule. However, without knowing the absolute expression in proteomics dataset, the summation rule will result in a compressed expression coefficient that reflects a 1.5-fold increase ((1+2)/(1+1) = 1.5).

For each of the 156 reactions whose flux and expression data are both available, we calculated the Pearson correlation coefficient (*PCC*) of the relationship between *relative expression of reaction* and flux using the 25 conditions as 25 datapoints. The *p-value* of each correlation was calculated based on a two-tailed hypothesis test. The correlation coefficients and *p-values* were calculated using the *corr* function in MATLAB 2019a. To adjust *p-values* for multiple testing, the *mafdr* function of MATLAB was used with the '*BHFDR'* method (Benjamini and Hochberg (BH) False Discovery Rate (FDR) correction). The resulting FDR values were used to evaluate the correlation. A correlation was considered significant if the FDR is less than or equal to 0.05.

*Analysis of pathway-level coexpression*

We manually defined 20 metabolic pathways that consist of at least three connected reactions for which flux measurements were available, and for which the enzyme levels corresponding to at least two reactions were available (Fig. EV 1e). To calculate pathway-level coexpression (Fig. 1e), we derived the *PCCs* of *relative expression of reaction* (over the 25 conditions) for every pairwise combination of reactions in a defined pathway and took the median value of these *PCCs*. Similarly, we defined the pathway-level flux-expression correlation as the median of the flux-expression *PCCs* (see above) for all reactions in a pathway.

To define coexpression patterns (Fig. 1f), we used the median of *relative expression of reaction* over pathway reactions, since median is robust to outlier reactions that do not coexpress with the majority of the pathway. For each one of the 20 pathways, we obtained 25 median levels that correspond to the 25 conditions. This 25-element vector was referred to as the pathway-level coexpression pattern.

Next, we calculated the *delta PCC* for reactions and pathways (Fig. 1f). To calculate the *delta PCC* for each reaction, we first correlated its relative flux over the 25 conditions with the coexpression pattern of the pathway that the reaction belongs to. Then, the original flux-expression PCC was subtracted from this PCC to derive *delta PCC*. The *delta PCC* of a pathway was defined as the median of *delta PCC* values for all reactions in the pathway.

*Analysis of pairwise cross-informing rate*

To calculate the cross-informing rate, all connected reaction pairs with measured flux and/or protein levels were collected. For instance, to calculate the flux-flux cross-informing rate, we collected every connected reaction pair with fluxes determined for both reactions. We calculated *PCC* and *p-values* for each collected pair using MATLAB, following methods described above. Reaction pairs with FDR less than 0.05 and *PCC* greater than 0 were deemed *cross-informed*. The cross-informing rate was defined as the proportion of cross-informed pairs in a given set of pairs. The sets of pairs were determined based on the network degrees of bridging metabolites, i.e., metabolites that connect the paired reactions. The network degree was defined as the total number of reactions that produce or consume a metabolite. Notably, reversible reactions contribute to the count of the network degrees of associated metabolites twice, since they can both produce and consume these metabolites. If a reaction pair was connected by more than one metabolite, the minimum network degree was used to define the bridging metabolite degree.

Importantly, due to the limited number of data points (i.e., only 232 reactions with measured fluxes), we grouped the pairs with approximate bridging metabolite degrees in the calculation of cross-informing rate. Specifically, for the calculation of flux-expression and flux-flux cross-informing rate, pairs with approximate bridging metabolite degrees were grouped until every group contained at least 20 pairs. After grouping, the bridging metabolite degree of each group was defined as the average degree of all pairs in the group. Each group produced a single data point in Fig. 2b in which the x-axis refers to the average bridging metabolite degree of pairs in the group and y-axis refers to the proportion of *cross-informed* reaction pairs. The grouping procedure is important to ensure that data points in Fig. 2b were calculated with similar sample size (~20 reaction pairs for each data point).

For the calculation of expression-expression cross-informing rate, the same grouping procedure was performed until every group contained at least 60 reaction pairs. The reason of using a higher threshold than 20 is that expression levels were available for more reactions than for flux data. The choice of sample size (60 and 20) was empirically made to best equalize the sample size of groups within a dataset (flux-flux, flux-expression, or expression-expression), while keeping the distribution of average degrees consistent between datasets.

*Mathematical formulation of Flux Potential Analysis (FPA)*

FPA is a specific flux balance analysis (FBA) problem that calculates the maximum flux potential (FP) of a target reaction (k) under certain constraints that address relative expression of reactions and their distance from the target (Equations S3-S6) [2]. The mathematical formulation of FPA is briefly summarized as follows. First, the metabolic model is converted to an irreversible model, where reversible reactions are split into forward and reverse reactions and all reactions carry non-negative flux. Second, as in any FBA problem [5], the steady-state assumption is imposed using Equation S4, where $s_{ij}$ indicates stoichiometric coefficient of metabolite *i* in reaction *j*, and $M_{all}$ and $R_{all}$ denote the set of all metabolites and reactions, respectively. Third, reaction fluxes are constrained between their upper and lower boundaries (Equations S5). Fourth, a weighted sum of flux is constrained to be less than or equal to a constant (Equation S6), which is referred to as ***flux allowance*** ($a$). Finally, the flux ($v$) of the target reaction is selected as the objective function, which is maximized to find FP as the objective value (Equation S3). Since the flux allowance is a constant, reactions with smaller weight coefficients ($w$) in the weighted sum (Equation S6) are more likely to carry larger flux to maximize the flux left for the target reaction. More detailed description of FPA could be found in the original study [2]. The enhanced flux potential analysis (eFPA) algorithm stays the same as original FPA except for the new distance decay functions and the use of weighted metabolic distance instead of naïve metabolic distance (see below).

$$FP_k = v_k^{max} = \max(v_k) \quad \text{S3}$$

$$s.t.$$

$$\sum_{j \in R_{all}} s_{ij} v_j = 0; i \in M_{all} \quad \text{S4}$$

$$0 \le v_i \le v_i^{UB}; i \in R_{all} \quad \text{S5}$$

$$\sum_{i \in R_{all}} w_i v_i \leq a, a = 1$$ S6

*Weight coefficients and distance decay function*

The key component of FPA is the calculation of the ***weight coefficients*** (*w*). Weight coefficient is defined as the product of gene expression coefficient ($c$), which is the reciprocal of the normalized expression level of a reaction, and the ***distance decay function***, which is a function of metabolic distance ($d$) (Equation S7). In our yeast analysis, the normalized expression level of a reaction is identical to the ***relative expression of reaction*** described in the previous section for correlation analysis. For reactions whose enzymes are not measured or only partially measured, a default gene expression coefficient ($c$) of 1 will be assigned to all conditions. The distance decay function represents the decrease of the influence of network reactions (enzyme reach) on FP as their distance from the target reaction increases. For instance, when estimating the flux potential of a reaction in the valine degradation pathway, one would not expect TCA cycle flux to be informative necessarily. In the original FPA, a power-law decay function (Equation S8) was proposed, in which gene expression coefficient is divided by its metabolic distance (plus one) to the power of ***distance order*** *(m).* A distance order of 1.5 was used for modeling *C. elegans* tissue metabolism [2]. As a result, for a reaction that is 10 reactions away from the target (*d = 10*), the weight in Equation S7 would be lowered by around 36 folds due to the second term, which would allow this reaction to take large flux values without significantly affecting FP calculations even if its expression coefficient is high (i.e., reaction is poorly expressed).

$$w_i = c_i \times f(d_{ik}), \;\; c_i = \frac{1}{normalized\ expression\ level\ of\ reaction\ i}$$ S7

$$f(d_{ik}) = \frac{1}{(1 + d_{ik})^m}$$ S8

In eFPA, we developed two new distance decay functions which we refer to as ***binary decay function*** (Equation S9) and ***exponential decay function*** (Equation S10)( Fig. EV 2c). In the binary decay function, a binary filtering was used so that the weight coefficient is not decayed if the distance of a reaction to the target is less than a given threshold, *n,* hereafter referred to as ***distance boundary***. In contrast, the weight coefficient is decayed to zero when the distance is greater than this distance boundary. For the exponential decay function (Equation S10), we designed a formula that represents an inverted S-curve (Fig. EV 2c). Compared with the binary decay function, this formula allows a smoother transition around *n*, which we still refer to as the distance boundary, before the value of function decays to zero.

$$f(d_{ik}) = \begin{cases} 1, \; d_{ik} \leq n \\ 0, \; d_{ik} > n \end{cases}$$ S9

$$f(d_{ik}) = \frac{1}{1 + 2^{d_{ik} - n}}$$ S10

*Weighted metabolic distance*

To integrate the bridging metabolite degree (Fig. 2a) with eFPA, we weighted the metabolic distance between two adjacent reactions based on the number of connections to the metabolite that connects them (Fig. EV 2b), such that the distance between reactions connected by hub metabolites is upscaled. Thus, the distance between a pair of reactions can be greater than the original distance that is the number of reactions between them plus one. We named the new distance measure as ***weighted metabolic distance***.

To calculate weighted metabolic distances, the metabolic network was converted to a directed graph where nodes represent reactions and edges represent the connectivity between reactions in the metabolic network. A directional edge was built between two reactions if (any of) the first reaction's product(s) could be used as the reactant of the second reaction. The reversible reactions in the network were split into separate reactions (forward and reverse direction) as two different nodes in the graph. In the calculation of the naïve (unweighted) distance, all the edges have uniform length (which is one), meaning each step in the path contributes to one unit of metabolic distance. However, for weighted distance, the length of edges is proportional to the network degree of the bridging metabolite (Fig. EV 2b; see also above**)**. We defined the length of edge between two reactions in weighted distance as ***degree of bridging metabolite/4***, where the division by 4 is to normalize the weighted distance to the same scale as the unweighted distance. After this normalization, the weighted distance is 1 for two adjacent reversible reactions in a linear pathway, which is equal to their unweighted distance. Together, the metabolic network was converted to a *weighted directed graph*, where nodes were (irreversible) reactions and edges were weighted metabolic distances between connected reactions (nodes).

The minimal weighted distance between any two reactions in the graph was calculated using the metabolic distance algorithm we have previously developed [2] with slight modifications. The original algorithm added a distance of 1 unit for every reaction that is traversed when going from one reaction in a pair to the other. In the new algorithm, the added distance per step is the edge length of the weighted directed graph, which is dependent on the bridging metabolite degree. Our distance tool is similar to Dijkstra's algorithm [6], one of the canonical algorithms for calculating shortest path in a graph. However, the shortest path cannot be guaranteed for all reaction pairs in a metabolic network due to reaction reversibility. The forward and reverse direction of an originally reversible reaction is represented by two individual reactions (nodes) in the converted graph (see above). Such pairs may coexist in a path that connects two distant reactions (i.e., the same reaction may be traversed twice, once in each direction), in which case the shortest distance path does not make a valid flux route. Mathematically, this forms a problem called *shortest path avoiding forbidden pairs* (PAFP) [7]. This problem has been shown to be NP-hard and canonical algorithms such as Dijkstra's algorithm cannot effectively solve it. Our new algorithm follows the same strategy as in the original [2] to cope with this problem. In brief, if both the forward and reverse forms of an originally reversible reaction coexist in a calculated shortest path, one of these nodes is removed from the network, and metabolic distance is recalculated.

This process may need to be repeated several times until a valid shortest path is found. We have previously shown that the copresence of both directions of a reversible reaction in the shortest paths is rare in metabolic networks [2] and confirmed the same observation in the weighted distance calculation (<1% for both yeast and human networks). Furthermore, in all cases that we have manually checked, the corrected shortest path produced by the iterative algorithm described above was truly the shortest path. Thus, PAFP had negligible effect on our analyses, if any.

Finally, the following metabolites were considered as by-products and were excluded from distance calculation, i.e., two reactions are not considered connected if they are only connected by the by-product metabolites [2]: carbon dioxide, AMP, NADP(+), NADPH, diphosphate, oxygen, NADH, NAD, phosphate, ADP, coenzyme A, ATP, H2O, H+, GTP, GDP, (R)-carnitine, FAD and FADH2.

*eFPA setup for SIMMER dataset*

The consensus yeast metabolic model was slightly modified and constrained before running eFPA. Four reactions were added to enable the secretion of orotate, enzymatic production of polyphosphate and energy consumption by non-growth-associated maintenance (NGAM), as described previously [1] (Table EV5). All possibly available nutrients (in the culturing media) were made freely exchangeable by setting the lower boundary of pertaining exchange reactions to -1000. These nutrients are phosphate, glucose, ammonium, uracil and leucine (Table EV5). No NGAM was imposed (lower boundary to 0) during eFPA. This constrained model was used in all eFPA analyses.

As in our previous analysis on *C. elegans* tissue metabolism [2], we set the expression coefficients ($c_i$) of all exchange reactions to zero (unless specified otherwise), in order not to penalize nutrient exchange twice (the other penalization is by the transport between extracellular space and cytosol). To account for media composition during eFPA, we modified nutrient exchange reactions in two ways. First, to address the absence of nutrients, we blocked the uptake of unavailable nutrients in each condition. Specifically, uracil and leucine uptakes were blocked in the eFPA of phosphate-limiting, carbon-limiting and nitrogen-limiting conditions (15 conditions in total); uracil uptake was blocked in leucine-limiting conditions (5 conditions); and leucine was blocked in uracil-limiting conditions (5 conditions). Second, to address the low abundance of nutrients (e.g., glucose concentration is over 20-fold lower in carbon-limiting conditions), we set an arbitrarily large gene expression coefficient ($c_i = 10$, Equation S7) on the pertaining exchange reactions. These included: (1) glucose exchange reaction (r_1714) for all carbon-limiting conditions (5 conditions); (2) phosphate exchange reaction (r_2005) for all phosphate-limiting conditions (5 conditions); and (3) ammonium exchange reaction (r_1654) for all nitrogen-limiting conditions (5 conditions).

*Running eFPA for SIMMER dataset*

eFPA was performed by a modified version of the generic eFPA function (https://github.com/WalhoutLab/eFPA) to enable highly parallel computation in a computer cluster. Solver parameters were set as previously described [2]. The distance boundary parameter was changed according to the question of interest as indicated in the text. All 232 reactions (Table EV2) with determined fluxes in SIMMER dataset were analyzed with eFPA.

*Correlation analysis of eFPA results and measured fluxes*

The direct output of eFPA is the flux potential values which were converted to relative flux potentials (rFP) as previously described [2]. This conversion includes dividing FP values with that of the super condition, a hypothetical condition where all enzymes are expressed at the highest level and which therefore yields a theoretical maximum FP for every reaction [2]. The rFP values were used to analyze correlation with measured fluxes. To evaluate the significance of correlations, in addition to the FDR threshold (0.05) used in the correlation of reaction level expression and fluxes, we applied a filter that removed reactions with low variation in rFP. Specifically, we required that the difference between the largest and smallest rFP (across 25 conditions) for a reaction be at least 0.2, an arbitrary threshold that defines significant variation above potential noise in experimental data.

*Titration of the distance boundary*

In Fig. 2d and Fig. EV 3, we systematically titrated the distance boundary and correlated the rFP values with measured fluxes. The distance boundary parameter was titrated from 0 to 40 with an interval of 0.5, resulting in 81 independent sets of predictions. Each prediction set was correlated to the fluxes to calculate *PCC* and FDR individually. The binary decay function was used for this analysis. A reaction was considered predicted with eFPA and was included in Fig. EV 3 if it showed significant correlation (*PCC* > 0, FDR < 0.05, min-max range > 0.2) in any of the 81 prediction sets.

*Calculation of effective distance boundary*

In eFPA, distance boundary parameter is measured in the scale of weighted metabolic distance. To relate this parameter to the actual length of the integrated pathway, we further converted it to an interpretable metabolic distance from the target (i.e., the maximum distance of integrated reactions to the target) that was used for data visualization in Fig. 2d and Fig. EV3.

To do the conversion, we first constructed a distance conversion table that mapped the weighted distance boundaries to the scale of naïve distance. For each of the 81 weighted distance boundary points used in titration (i.e., from 0 to 40 with uniform intervals of 0.5), we took all reactions within the weighted distance boundary of the target, and calculated the maximum naïve distance of these reactions to the target, which we call the *effective metabolic distance boundary*. This procedure was performed for each of the 232 reactions and for the 81 distance boundary parameters to generate the conversion table.

Next, we used the distance conversion table to transform the original 232-by-81 rFP-flux *PCC* matrix with the weighted distance boundaries as rows into a 232-by-41 matrix with the effective boundaries as rows. For each target reaction, given an effective distance boundary, we identified *PCC* values with matching effective distance in the conversion table and took the maximum of these *PCCs* to define the converted *PCC* in the transformed matrix. When there was no match in the conversion table (i.e., all effective distances were less or greater than the desired value for the target reaction), the converted *PCC* for the nearest smaller effective distance boundary was used. Finally, the resulting 232-by-41 *PCC* matrix was row-wise normalized by dividing the maximum *PCC* and then visualized in the heatmap of Fig. EV3 and Fig. 2d.

*Deciphering prediction mechanisms of eFPA*

The mechanisms that drove an eFPA prediction can be deciphered from the flux distribution that maximized FP (Equation S3). Since total flux allowance has to be equal to or less than the flux allowance ($a$) (Equation S6), most reaction fluxes tend to be minimized unless the high level of a particular flux is required to fulfill the flux maximization of the target reaction. For those required fluxes, their weight coefficients ($w$) therefore significantly impacted the FP of the target (Equation S6). Thus, the reactions with significant flux allowance contribution (defined by the product of its weight coefficients and flux) are the major contributors to the eFPA prediction, and their *relative expression of reaction* collectively determines the FP of the target reaction. We calculated the flux allowance contribution of every reaction by multiplying the weight coefficient with its flux in the resulting flux distribution of each eFPA calculation. The results were visualized in Appendix Fig. S2a.

*Randomization test of eFPA*

To assess the statistical significance of eFPA modeling, we shuffled the rows (reaction labels) of the *relative expression of reaction* matrix (658 reactions by 25 conditions), i.e., randomized the association between the expression levels and reaction labels. After shuffling, eFPA was performed with randomized data following the same procedure as described above. This randomization was performed for 1000 times. It is noteworthy that we were unable to perform a greater scale of randomization due to the overwhelming computational demand.

*eFPA on the drainage flux of biomass precursors*

To construct the objective function that represents the drainage flux of a biomass precursor, we added a demand reaction that could consume the target metabolite. To consistently analyze biomass precursors with variable network degrees, we assigned a uniform weighted metabolic distance of 1 between the added demand reaction and reactions that produce the corresponding biomass precursor regardless of the bridging metabolite degree. Other configurations remained the same as in regular eFPA with the 232 reactions. The eFPA results of biomass precursor draining fluxes were visualized following the same effective distance boundary conversion and normalizations as stated above for Fig. EV3.

*Other analyses for SIMMER dataset*

In Fig. 2f,g, we compared eFPA with predictions made from simple average of local expression, as well as from target-masked enzyme levels. For calculating the local expression average, the average value of the *relative expression of reaction* for all qualified reactions was calculated. The qualified reactions were defined as reactions whose (i) metabolic distance is less or equal to the specified distance boundary, and (ii) the expression level was determined. The resulting average values were then correlated with flux to calculate *PCC* and FDR using the MATLAB functions stated above.

For the masking of target expression (Fig. 2g), we set the gene expression coefficient of the target reaction to 1 in all conditions. This is mathematically equivalent to the scenario in which the expression of the target reaction is not measured.

<u>eFPA Analysis on Human Tissue Dataset</u>

*Processing of tissue expression data*

Quantitative proteomics and transcriptomics of human tissues were obtained from the supplementary materials of Jiang et al. [8], which include RNA and protein levels of more than 12,000 genes across 32 normal human tissues quantified based on 201 individual primary samples. Our tissue metabolism analyses were performed at the level of 32 tissues by using the *tissue median* provided in the referred study. We rescaled both RNA and protein tissue medians to make them suitable for system-level modeling.

For RNA-seq data, we converted the tissue median log2TPM to raw TPM (Equation S11). The raw TPM was used as the input for downstream analysis and was referred as ***RNA tissue median***.

$$RNA\ tissue\ median = 2^{tissue\_median\_\log_2 TPM} \quad \text{S11}$$

To robustly model tissue specificity in eFPA, we further constructed a noise-robust metric of tissue median expression that combines the Tissue Specificity score (TS score), a Z-score-like metric representing the deviation from the population mean for each tissue [8], with the tissue median. A shortage of modeling based on tissue median instead of individual samples is the loss of variance information. For example, the expression of a gene may be highly variable due to genetic heterogeneity in the population. In this case, the median expressions of different tissues may also vary largely, however, this variation is not necessarily an indication of tissue heterogeneity. To avoid misinterpretation led by high variation instead of real tissue specificity, we combined the tissue median with the TS score that informs the variance level. Since TS score describes how much a tissue median deviates from the population mean, we could convert it into a *p-value* that indicates the probability of the tissue median being greater or less than the population mean (Equation S12, where the cumulative distribution function normcdf returns values between 0.5 and 1 since the absolute value of TS score is used). Importantly, one minus the *p*-value indicates the confidence level of tissue specificity. To combine this confidence level estimate with the tissue median, we used it as a weight coefficient to compress the log2-Fold-Change (log2FC) of each tissue median compared with the population mean that was calculated during TS score modeling in the referred study [8] (Equation S13, where log2FC is represented by the difference between log2TPM values). If a tissue median only presents insignificant deviations from the population mean because of the expression level itself (log2TPM) being close to the mean or because of a large p-value, this formula pushes the calculated metric toward the value of the population mean (as TPM) by compressing the second power term to zero. We termed this metric as ***compressed RNA tissue median***.

$$P(TS\ score) = 2 \times (1 - normcdf(abs(TS\ score)))$$
$$normcdf: cumulative\ distribution\ function\ of\ N(0,1) \quad \text{S12}$$

$$compressed\ RNA\ tissue\ median = 2^{population\_mean\_\log_2 TPM} \times 2^{(1-P(RNA\ TS\ score)) \times (tissue\_median\_\log_2 TPM - population\_mean\_\log_2 TPM)} \quad S13$$

For proteomics data, the reported tissue median was the median log2FC against a meta-reference sample. We recentered this value to the corresponding protein population mean estimates [8], by subtracting the population mean, so that genes with no tissue-enriched expression will have a log2FC around zero. We also positioned missing abundance values, an artifact of proteomics [8], at this center to keep them in line with the null hypothesis (not tissue enriched). In addition, we rescaled the recentered log2FC with estimated absolute abundance to facilitate eFPA modeling. As described in the yeast eFPA above, proteomics data, as a FC-based relative expression measurement, poses challenges in the modeling of complex GPR involving isozymes (i.e., when the expression levels of individual genes need to be summed up to reach a reaction level result). To address this issue, we used the matched RNA-seq data to rescale the proteomics data such that absolute abundance of each protein is roughly represented. We used the population mean of absolute RNA abundance as a quantitative proxy of the population mean of absolute protein abundance. Therefore, the absolute abundance of protein could be estimated by multiplying the RNA population mean with the proteomics-derived FC. Finally, we applied the same idea used for compressing the RNA data (see above) to obtain the ***compressed protein tissue median*** (Equation S14).

$$compressed\ protein\ tissue\ median = 2^{population\_mean\_\log_2 TPM} \times 2^{(1-P(protein\ TS\ score)) \times (tissue\_median\_\log_2 FC^{vs.\ meta\ ref} - population\_mean\_\log_2 FC^{vs.\ meta\ ref})} \quad S14$$

*Processing of tissue-enriched metabolite set*

The tissue enriched metabolite sets were obtained from MetaboAnalystR software (version 4.93, www.metaboanalyst.ca) [9]. There were 73 tissue-enriched or subcellular-enriched metabolite sets in the original data table. Tissue labels in MetaboAnalystR metabolite set were manually matched with the 32 tissues measured in Jiang *et al.* [8] (Table EV7). A total of 17 distinct tissue types were assigned to one or more matched tissues in both MetaboAnalystR and Jiang *et al*. study. When more than one of the tissues are matched with the same tissue type (e.g., Adrenal Cortex, Adrenal Gland and Adrenal Medulla are all assigned to Adrenal Gland), their metabolite sets were merged. Similar merging process was also done for the eFPA predictions (see below). Metabolites in the MetaboAnalystR set were matched with those in the human metabolic model by their HMDB ID or manually when needed. In the end, 331 metabolites from the human metabolic model were labeled with an enrichment in at least one of the 17 tissue types, resulting in 827 tissue-metabolite pairs in total (Table EV8). Of the 17 tissue types, 8 major tissue types have more than 50 tissue-enriched metabolites (Brain, Skin, Liver, Intestine, Muscle, Pancreas, Prostate, Spleen), making them relatively comprehensive references for benchmarking. The remaining 9 tissue types have fewer tissue-enriched metabolite assigned, ranging from 1 (Colon) to 37 (Testis).

*Human metabolic network model*

The newest consensus human metabolic model, Human1 (version 1.5.0), was downloaded from metabolicatlas.org [10]. The model was first converted to COBRA-format model by *ravenCobraWrapper* function in RAVEN toolbox [11]. Next, we made the following modifications to the model to make it suitable for modeling tissue metabolism.

First, large coefficients in the stoichiometry matrix were reduced to avoid numerical stability issues during Mixed-Integer Linear Programming (MILP). In the human model, large coefficients typically appear in reactions that assemble high-molecular-weight molecules such as antigen. We rescaled these reactions, while maintaining mass balance, to make sure the maximum coefficient in the stoichiometry (S) matrix is less or equal to 100. A hypothetical example is provided in Equation S15.

$$\begin{aligned}&hypothetical\ original\ formula:\\&\qquad 1000A + B = C\\&rescaled\ formula:\\&\qquad 100A + 0.1B = 0.1C\end{aligned} \qquad \text{S15}$$

Second, the stoichiometry of reactions that carry very small flux was also adjusted to avoid numerically zero fluxes that are actually significant. Low-flux reactions typically involve micronutrients such as vitamins and carry fluxes as low as $10^{-7}$ under regular boundary constraints. We adjusted the stoichiometric coefficients of these reactions proportionally (see a hypothetical example in Equation S16) to make sure their maximum flux is always greater than 1e-4 in flux variability analysis (FVA), unless they carry no significant flux.

$$\begin{aligned}&hypothetical\ original\ formula\ (Vmax = 10^{-7})\\&\qquad A + B = C\\&rescaled\ formula\ (Vmax = 10^{-4})\\&\qquad 0.001A + 0.001B = 0.001C\end{aligned} \qquad \text{S16}$$

Third, to model the differential availability of nutrients in blood stream, we divided the nutrients into two groups, major and side nutrients, and controlled their uptake via a set of uptake reactions that replaced corresponding exchange reactions of the original model. In total, 38 metabolites were assigned as major nutrients (Table EV9) because they are (1) abundant in serum according to serum metabolomics database [12]; or (2) lipid carrier protein complexes such as HDL cholesterol; or (3) amino acids. Other nutrients were considered as side nutrients. In addition, 10 inorganic metabolites (e.g., oxygen) were excluded from major and side nutrient sets and were made freely exchangeable (i.e., lower boundary fluxes were set at -1000) (Table EV9). We controlled the mass influx of each type of imported nutrient using a flux balance method that we developed previously [2]. This method employs specialized uptake reactions that import a nutrient and a pseudo-metabolite, and exchange reactions that drain the pseudo-metabolite. Briefly, for each major or side nutrient, a new uptake reaction was added to import the nutrient into the system. Each uptake reaction produces 100 mg of the target nutrient, along with 1 unit (mmol) corresponding pseudo-metabolite (Equation S17). Two pseudo-metabolites were added to the model, *majorNutr* and *sideNutr.* They are drained by two corresponding exchange reactions. They serve as proxies of the mass influx of major and side nutrients in FBA, respectively. For example, 1 unit of flux (mmol/h/gDW) in the uptake reaction of glucose (Equation S17) produces 1 mmol *majorNutr* and 100 mg glucose (~0.55 mmol). The *majorNutr* is then drained by the exchange reaction of *majorNutr*, EX_majorNutr, with a coefficient of 10 (Equation S18). Thus, one unit flux of EX_majorNutr allows 1 gram of nutrient to be imported. Similarly, the total influx of side nutrients is controlled by the exchange reactions of *sideNutr*. During the reconstruction of uptake reactions, we encountered extremely low metabolite coefficients (i.e., $<10^{-5}$ instead of 0.555069 in Equation S17) for polymeric metabolites associated with extremely high molecular weights in the human model. We adjusted the coefficients of these reactions as we did with reactions that have large coefficients and low-fluxes (see above, Equations S15 and S16), except that the values of coefficients were proportionally increased instead of decreasing. This adjustment meant the individual metabolite uptake was >100mg per unit flux of the adjusted uptake reaction, but the equivalence of 1 unit flux of EX_majorNutr or EX_sideNutr to 1 g material did not change. In the end, the stoichiometric coefficients of the model were kept in the range $10^{-5}$ and 100 while mass balances were properly maintained. After reconstructing the uptake reactions, the original exchange reactions of side and major nutrients were constrained to [0, 1000], only allowing secretion. And the default boundary of uptake reactions were set at [-1000, 0], only allowing uptake. The total uptake fluxes are eventually controlled by the two exchange reactions of pseudo-metabolites. Their constraints were defined based on specific modeling purposes. The default was set to [0, 1000] that allows for unlimited uptake of both major and side nutrients. This set up was used to control nutrient availability when building human tissue metabolic networks as will be explained below.

$$\begin{array}{l} example\ of\ uptake\ reactions: glucose\ (m01965s[s])\ uptake \\ \quad 0.555069\, m01965s[s] + majorNutr\ <=> \end{array} \qquad \text{S17}$$

$$\begin{array}{l} exchange\ reaction\ of\ majorNutr\ (\text{EX\_majorNutr}): \\ \quad 10\, majorNutr\ -> \end{array} \qquad \text{S18}$$

Forth, the Gene-Protein-Reaction association (GPR) of pyruvate dehydrogenase reactions was modified to “ENSG00000091140 and ENSG00000110435 and (ENSG00000131828 or ENSG00000163114) and ENSG00000150768 and ENSG00000168291” (Hao Wang *et al*. [10], personal communication) to correct an annotation error.

This final modified model was used in both tissue network building and eFPA analysis.

*Building human tissue networks*

To comprehensively model the tissue metabolism in humans we followed the modeling pipeline MERGE, which we previously developed for modeling *C. elegans* tissue metabolism [2]. In brief, two stages of modeling were performed in a sequential manner. In the first step, a semi-quantitative modeling of on/off status and direction of reaction fluxes was performed using iMAT++ algorithm [2]. This step is to globally fit the distinct expression levels of enzymes (see below) in tissues to the metabolic network, to derive (1) a flux distribution that we named as Optimized Flux Distribution (OFD) and that can be used to assign the flux directions for reversible reactions; and (2) a tissue-specific metabolic network in which inactive reactions with zero flux were removed from the network. After iMAT++ modeling, the eFPA analysis was performed on the derived tissue networks. Using a tissue network instead of the naïve network (i.e., all reactions included) provides a more realistic network context for the eFPA analysis. In addition, the rFP predictions generated from eFPA was overlayed with OFD to derive high-confidence predictions with reaction directionality. This step is important for the interpretation of eFPA predictions of reversible reactions that show high flux potential in both directions, such that the direction predicted by iMAT++ is chosen as the most likely metabolic function with high potential. Together, this tissue modeling pipeline provides a comprehensive collection of high-quality predictions about tissue-enriched metabolic fluxes.

Two sets of tissue networks were built for the 32 tissues based on either proteomics data (protein-based tissue network; PTN) or RNA-seq data (RNA-based tissue network; RTN). Parameters, settings and modifications are summarized below and further details about the MERGE modeling pipeline can be found at our previous publication [2].

*iMAT++: expression categories*

For building RTN, metabolic genes were categorized by published procedures [2] with slight modifications. In brief, 32 tissue median log2TPM profiles (*RNA tissue median*) were used as the input data. A tri-modal mixed Gaussian curve was fitted to the aggregated histogram of all log2TPM data using the *fitgmdist* function in MATLAB. The fitted means of the three components (log2TPM= -7.2, 0.64, and 4.2) were used as the thresholds to classify rarely (log2TPM < -7.2), lowly (-7.2 <= log2TPM < 0.64), moderately (0.64 <= log2TPM < 4.2) and highly (log2TPM >= 4.2) expressed genes. This produced the primary gene categories. To further address the tissue-enriched genes, we refined the primary gene categories based on the TS score of each gene. We reconsidered the gene categories for genes whose TS score is higher than 2.5 or lower than -2.5 (in a tissue). For those higher than 2.5, the following rules were used to refine the categories: (1) originally lowly expressed genes were moved to moderately expressed category, to not avoid flux on tissue enriched genes; (2) originally moderately expressed genes were moved to highly expressed category to encourage flux on genes that are both expressed and tissue-enriched; (3) gene category was not changed if the gene is originally in rarely or highly expressed category. We didn't find any gene that shows TS score greater than 2.5 in rarely expressed category. For those with TS lower than -2.5, we only moved originally moderately expressed genes to lowly expressed category, to address their tissue-depleted feature. After refinement, the final gene categories were used in iMAT++.

For building PTN, we used a hybrid approach to derive the gene categories. Since the proteomics data lacks absolute expression information, we used the primary gene categories derived by RNA-seq data, thus using the absolute abundance of RNA as a proxy of the absolute abundance of protein. Next, we refined the gene categories based on relative expression as with the RTN categorization. The only exception is that if a protein was shown to be tissue-enriched (TS > 2.5) and was in rarely expressed category in the primary gene categories, we moved it to the moderately expressed category, since proteomics data indicate that these enzymes were expressed regardless of their extremely low mRNA levels. There are four such genes in total (ENSG00000205186, ENSG00000144035, ENSG00000182591, ENSG00000184210).

*iMAT++: flux thresholds*

iMAT++ uses a set of user-supplied thresholds (hereafter epsilons) to define significant flux when forcing fluxes on highly expressed reactions/genes [2]. The default epsilon value was set at 0.01. As in our previous work, we lowered the epsilon for reactions who cannot carry a flux of 0.01 in normal conditions. Specifically, we first performed FVA while allowing 10 units (10 grams) major nutrient exchange flux and 1 unit (1 gram) side nutrient exchange flux. Then, the epsilon of any reaction whose maximum flux is lower than 0.01 was adjusted to 10% of this maximum flux.

*iMAT++: algorithm modifications*

The original iMAT++ algorithm was used with two minor modifications.

Firstly, during the inspection of the above-mentioned gene categories, we found that some electron transport chain (ETC) genes were lowly or rarely expressed in most if not all tissues. However, the vast majority of ETC components were actively expressed and should be functioning in all tissues. This discrepancy may be due to wrong GPR annotations, detection problems, or the thresholding approach not being suitable for some ETC genes. Since oxidative phosphorylation is the core energy production pathway in most tissues, blocking ETC flux may have a global impact on all FBA-based procedures. As a remedy, we forced flux through ETC during iMAT++ integrations. Specifically, we added a new functionality to iMAT++ algorithm where we could specify a core reaction set that is treated as highly expressed regardless of the categories of the associated genes. Notably, the core reaction idea was already used and implemented in the original iMAT algorithm, which is the predecessor of iMAT++ [13,14]. The core reactions used in this study are CYOOm3i, HMR_6911, HMR_6912, HMR_6914, HMR_6916, HMR_6918, HMR_6921.

Secondly, to model the differential nutrient availability in blood circulation, we decided to minimize the usage of side nutrients in the iMAT++ integration. To achieve this while keeping minimal interference with the fitting of gene categories, we replaced the original total flux minimizations with a two-step flux minimization. In the first step, we minimized the total side nutrient usage by minimizing the flux through EX_sideNutr reaction. Next, the maximum side nutrient usage was constrained to 110% of this minimized flux. Finally, the total flux was minimized given the 110% side usage cap in addition to other fitting constraints. This modification was applied to all flux minimizations in iMAT++ procedure including the calculations for Primary Flux Distribution (PFD), latent flux fitting, and OFD [2].

In other parameter settings, we allowed unlimited major nutrient uptake (i.e., upper boundary of EX_majorNutr was set to 1000) during iMAT++. The Speed Level was set to level 2 to balance the computation efficiency and MILP optimality [2]. Waste of major nutrient in OFD was assessed as a quality control of the integration, similar to the bacterial waste assessed in our previous *C. elegans* modeling [2]. Most tissues OFD had zero percent waste, which indicated the major nutrients in blood stream were used as major carbon and energy source in the metabolism.

*Building tissue networks based on FVA*

The second step of iMAT++ is to build the context-specific networks by Flux Variability Analysis (FVA). In the standard pipeline, FVA is performed on the MILP model constrained with all fitting constraints (see our previous work [2] for details). FVA measures the full solution space of the flux of each reaction, and thus we can obtain a set of inactive reactions that cannot carry significant flux given the input gene categories and flux constraints. The network with those inactive reactions removed is the context-specific network.

Large-scale FVA could be computationally intensive, especially with large models such as human 1. To alleviate this problem, we developed a faster computational equivalent of FVA to build the network, which we call Flux Accessibility Analysis (FAA). FAA derives the identical context-specific network as FVA but is twice as fast as FVA in speed. The reason for the improved speed is that FAA reforms the optimization problem of FVA to avoid extensive numerical optimization. Originally, the network building had two steps: first, a whole-network FVA is performed to obtain the upper and lower flux bound of each reaction. Second, the upper bound and lower bound of each reaction are compared with a user-defined significant flux threshold (i.e., epsilon) to determine if a reaction is inactive. The disadvantage of this strategy is that FVA often spends a lot of time on fine-tuning the boundaries to get the exact maximal and minimal flux, although it is easier to check if the boundary already passes the significant flux threshold, which is the only information needed for building the network (i.e., whether the reaction can carry flux). Therefore, we developed FAA, which uses an integer objective function to test if the target reaction could carry flux higher or equal to the significant flux threshold as shown in Equations S19 and S20, where the subscripts *f* and *r* denote forward and reverse directions of a reaction and *y* is a binary integer variable. The $\varepsilon_f$ and $\varepsilon_r$ values are the epsilons for the forward and reverse direction with $\varepsilon_r$ being a negative value. In addition to improving the computational speed, FAA also provided higher numerical precision since the relative MIP gap tolerance could be set to 1e-12 instead of 1e-4 used previously in speed level 2 FVA [2]. Having replaced FVA with FAA, we followed the network-building rules in our *C. elegans* modeling [2] to classify reactions into OFD (i.e., carrying flux in optimal flux distribution), ALT (carrying flux in alternative solution space) and SLNS (not carrying flux in solution space) status. Please refer to the source code (https://github.com/WalhoutLab/eFPA) and supplementary information for iMAT++ [2] for details.

$for\ a\ target\ reaction\ with\ flux\ \boldsymbol{v}\ and\ significant\ flux\ threshold\ \boldsymbol{\varepsilon}$

$$v_f \geq v^{LB} - y_f\left(v^{LB} - \varepsilon_f\right); y_f \in \{0,1\} \qquad \text{S1}$$

$$FAA_f^{obj} = \max\left(y_f\right) \qquad 9$$

$$v_r \leq v^{UB} - y_r(v^{UB} - \varepsilon_r); y_r \in \{0,1\}$$
$$FAA_r^{obj} = \max(y_r)$$ S2

0

The speed of iMAT++ based network building (OFD and FAA) of human tissue networks is ~1 hour per tissue in a 20-core lab server. The resulting 32 tissue networks were further used in eFPA.

*eFPA analysis: general setup*
We performed comprehensive eFPA with different tissue networks (PTN and RTN), expression datasets (proteomics, RNA-seq) and objectives (regular flux and boundary flux). We explain the procedures of each analysis in detail in the sections that follow.

eFPA was performed on the same model used in iMAT++ network building. As in the yeast modeling, we also set the expression coefficients to zero for all exchange reactions (both original exchanges and new uptake reactions). Nutrient uptake was unlimited by setting the upper bounds of major nutrient and side nutrient exchange to 1000. When not specified, the iMAT++ derived tissue networks were used in eFPA analysis. PTN and RTN were used in eFPA based on proteomics and RNA-seq data, respectively. For solver parameters, we set both *optTol* and *feasTol* to 1e-7 as a default. Default flux allowance ($a$) was 1. The FPA method works with the assumption that this flux allowance is limiting (see Equations S3-S6). However, when the target reaction has very low flux carrying capacity, the FPA solution may not use the entire allowance, in which case this assumption is violated and in extreme cases flux potentials may be uniform for all tissues regardless of gene expression coefficients. A condition that warrants flux allowance to be limiting is the maximum flux carrying capacity of the target (based on regular FVA) being greater than or equal to 1. We therefore separately calculated eFPA of all reactions with flux carrying capacity <1 using smaller flux allowance values as well as a lower *feasTol*. More specifically, the flux allowance value was always made lower than the maximum flux carrying capacity of the target and *feasTol* was set at 1e-9 to account for low fluxes encountered in these simulations. Finally, the decay function of eFPA algorithm in human tissue modeling was the exponential decay with a distance boundary of 6 (the local-pathway eFPA), and the distance measurement is the weighted metabolic distance.

*eFPA analysis: metabolic distance*
To calculate weighted metabolic distance, the following metabolites were considered as by-products and were excluded from distance calculation: $CO_2$, AMP, NADP+, NADPH, PPi, $O_2$, NADH, NAD+, Pi, ADP, CoA, ATP, $H_2O$, H+, GTP, GDP, Electron Transfer Flavoprotein Reduced, Electron Transfer Flavoprotein Oxidized, L-carnitine, FAD, $FADH_2$ and Na+. Other procedures stayed the same as for the yeast network.

*eFPA analysis: expression data*
We used compressed RNA and protein tissue medians (see above) as the input expression data for eFPA. In the figures and main text, we used a terminology regarding the input dataset for eFPA, which is explained in detail below:

- **Protein eFPA**: *compressed protein tissue median* was used as the input dataset. However, to make fair comparison with RNA eFPA, we only took the commonly detected genes in both *compressed RNA tissue median* and *compressed protein tissue median* datasets. In total, 12,121 genes were detected in both datasets, covering 2869/3627 (79%) genes in the Human 1 model.
- **RNA eFPA (common genes) or RNA eFPA**: *compressed RNA tissue median* was used as the input dataset. As with the protein eFPA, only the intersection genes were used in the modeling.
- **RNA eFPA (all genes)**: *compressed RNA tissue median* with all detected genes was used as the input dataset. Notably, the RNA-seq detected more genes than proteomics, so there were 19,273 genes detected, covering 3603/3627 (99%) genes in the Human 1 model.

When not specified (e.g., referred to as **eFPA**), the default dataset used was the *compressed protein tissue median* with only the genes overlapping with the RNA dataset (n=12,121).

*eFPA analysis: regular reactions*
We analyzed all regular reactions in human 1 model except for transport, exchange, and custom uptake reactions (see above). The computation was performed in Massachusetts Green High Performance Computing Center (GHPCC) with 512 cores. The eFPA of all internal regular reactions (7,248) and 32 tissues took less than 30 minutes.

*eFPA analysis: boundary reactions*
For boundary reactions, we performed eFPA on transporter reactions that take up or secrete metabolites from or into extracellular space, respectively. As previously done for transporter FPA [2], we applied the treatments to avoid shortcuts, such as the elimination of demand and sink reactions that drain or introduce the same metabolite as the target transport reaction. All transporter reactions that exchange metabolites between a cellular compartment and the extracellular compartment were analyzed (2,852 reactions).

*Linking boundary flux predictions to tissue-enriched metabolites*
eFPA produced an rFP matrix with columns representing the 32 tissues and rows the reactions (i.e., internal reactions and transporters). We next row-wise centered this matrix by subtracting the row median, so that we obtained the delta rFP from the median (ΔrFP). Thus, larger ΔrFP values indicate tissue enrichment.

Since the boundary flux may be a quantitative proxy for the tissue abundance of the corresponding metabolite, we used the maximum ΔrFP among the transporter reactions that import or export a target metabolite as the tissue enrichment score of this metabolite. This yielded predictions for 289/331 metabolites that we had reference data for (see above). These 289 metabolites accounted for 740/827 metabolite-tissue pairs that were annotated as tissue-enriched.

To generate the predictions only based on expression of target transporters (Fig. 4c), we performed original FPA with distance order of 100 and regular metabolic distance, which generates rFP only based on the expression of the target reaction [2]. This method gives identical results with using the *relative expression of reaction* as rFP. We used original FPA for convenience, as it was already built into our tissue modeling pipeline.

If a tissue type in the benchmark dataset is related to more than one tissue in eFPA, e.g. “Heart” is related to both “Heart Atrial” and “Heart Ventricle”, the maximum ΔrFP from the pertaining tissues was taken as the final ΔrFP of this tissue type. The final ΔrFPs for the 289 metabolites in the 17 annotated tissue types were used in further evaluations.

*Evaluating tissue-enriched metabolite predictions*
We performed statistical enrichment analyses to evaluate the tissue enriched metabolite predictions represented by ΔrFPs calculated for transportable metabolites. First, we considered the hypergeometric enrichment between the prediction and the reference set for metabolites in the reference set. To convert the ΔrFPs into binary calls of tissue enrichment (*enriched* or *not enriched*), we set a threshold on the ΔrFPs, where metabolite-tissue pairs with ΔrFPs higher than this cutoff were labeled as *tissue enriched*. We used hypergeometric test to check if the predicted *tissue enriched* metabolite-tissue pairs significantly overlap with the reference *tissue enriched* metabolite-tissue pairs. We varied the ΔrFP threshold to assess the significance of enrichment with different cutoffs.

Second, we considered the enrichment of high ΔrFP values in the known tissue-enriched metabolites. The distribution of ΔrFP values for all transportable metabolites in the network (n=1,379) represented a null distribution of non-tissue-enriched metabolites. We compared this overall distribution with the ΔrFP distribution of the 740 reference tissue-metabolite pairs using boxplots. We also tested whether the median ΔrFP of tissue-enriched metabolites is greater than the median of the background via rank-sum test.

*Clustering rFPs of internal reactions*
In Fig. 4d, we show the clustergram of ΔrFP of human tissue-enriched metabolism. The data processing for and the generation of this figure are summarized here. First, we filtered the ΔrFP predictions by a cutoff of 0.2. Only reactions whose ΔrFP were greater than 0.2 (tissue-enriched) or lower than -0.2 (tissue-depleted) in at least one of the 32 tissues were kept. We kept reactions that satisfied this criterion in either RNA-based prediction or protein-based prediction. Second, to address the cases when eFPA predicts tissue enrichment for both directions of a reversible reaction, we selected the eFPA predictions based on the relevant flux distribution (OFD) predicted by iMAT++ [2]. The reactions that passed these criteria were merged at the end. We termed these final filtered predictions *high-confidence predictions of human tissue metabolism* and generated the clustergram with them. The clustergram was generated with *clustergram* function in MATLAB with ‘cosine’ distance for both row and column clustering. These high-confident predictions were starred (i.e., labeled with “*”) on their reaction ID in Table EV6.

*Clustering tissue-enrichment of subsystems*

To facilitate the interpretation of tissue-enrichment of metabolism, we grouped the reaction-level predictions into subsystems so that the subsystem-level metabolic specialization in tissues can be visualized. To obtain the subsystem-level information, we first defined a reaction as tissue-enriched if the ∆rFP is greater than 0.2. Thus, each of the 32 tissues was assigned with a set of tissue-enriched reactions. Next, we generated a matrix where rows are subsystems and columns are the 32 tissues. The values in the matrix are number of tissue-enriched reactions assigned to each subsystem and tissue. Finally, for visualization, we row-wise normalized this matrix by dividing all values with the maximum, which yielded the relative tissue-enrichment for subsystems shown in the heatmap (Appendix Fig. S3).

### Supplementary Notes

Enhanced Flux Potential Analysis

Enhanced Flux Potential Analysis (eFPA) is a derivative of the flux potential analysis (FPA) algorithm that we previously developed [2]. Although the framework stays the same, critical redesigns were made in the eFPA. Here we introduce the concept of FPA and redesign of eFPA.

The goal of FPA is to predict the relative flux potential (rFP) for each metabolic reaction (defined as target reaction) under each condition based on gene expression data. The contribution of each non-target reaction to the rFP of the target reaction decreases with the network distance of the non-target to the target reaction following a distance decay function (Fig. EV2a). The process of integrating gene expression is formulated as an FBA problem whose maximized objective value is named *flux potential (FP)*. rFP is the normalized FP for which the FP is divided by the FP of the super condition, a hypothetical condition that provides the theoretical maximum FP of a reaction (see Methods). Thus, rFP is a value between zero and one. The key concept of this FBA problem is to convert the relative levels of enzyme expression into weight (penalty) coefficients such that the lower a relative expression is, the higher is the weight to penalize the flux through the corresponding enzyme. Therefore, a high rFP is expected only if the relative expressions of reactions surrounding the target within the flux route of the FBA solution are consistently high (farther reactions play a lesser role since their weights are always low due to distance decay). The mathematical details of FPA can be found in Supplementary Methods.

In eFPA, redesigns were made regarding metabolic distance measurement and the distance decay function. First, to integrate the idea of bridging metabolite degree (Fig. 2a), we redefined the metabolic distance using additional weight coefficients (Fig. EV2b). These weight coefficients increase the distance value between the target reaction and reactions that connect to this target by hub metabolites, thereby reducing the influence of such reactions in eFPA. We found that the new distance metric shows a more normal-like distribution with a greater mean compared to the original (referred to as *weighted distance*, Fig. EV2b). Secondly, we redesigned two distance decay functions, named the *binary decay* function (used in optimal-boundary eFPA) and the *exponential decay* function (used in local-pathway eFPA), both of which work with a single parameter *distance boundary* (Fig. EV2c). To precisely control the contribution of non-target reactions, i.e., to explicitly define the length of pathway being integrated, we used the binary decay function, in which the contribution of non-target reactions to the target reaction decays with the distance from the target by a Z-curve (Fig. EV2c), such that, the expression information within the specified distance boundary (represented by the upper plateau of the Z-curve) is integrated equally, while that beyond this boundary (the lower plateau) is filtered out. For better parameter robustness in flux prediction applications, we used the exponential decay function, in which the contribution decays with an S-curve that imposes a smoother transition around the boundary (Fig. EV2c). Together, eFPA performs controllable analysis of enzyme expressions in related pathway(s) to predict relative flux potentials of reactions of interest in a metabolic network.

Associating flux indicators with their target reactions

To visualize the association between an indicator enzyme and its target reactions that carry colinear fluxes with this indicator, we constructed a metric named co-correlation coefficient. The co-correlation coefficient is the product of *PCC* between the indicator's flux and target's flux and the *PCC* between the indicator's expression and target's flux. Thus, the co-correlation coefficient represents both the collinearity of the two fluxes and strength of correlation between the flux of the target and the expression of the indicator, and takes a high value only if both of these values are close to 1. In Fig. EV4, we visualized the co-correlation coefficient between the 46 correlated reactions (as potential indicators, columns of the heatmap) and their flux-colinear counterparts (collinearity > 0.8, rows of the heatmap). The big clusters in the heatmap were usually related to pathway-level coregulations such that multiple reactions in the pathway can serve as indicators (e.g., arginine biosynthesis) for the entire pathway. In comparison, the vertical strips (e.g., r_1022 vs. TCA reactions) revealed the single indicators for the colinear reaction sets, which were discussed as examples in Fig. 3b,c.

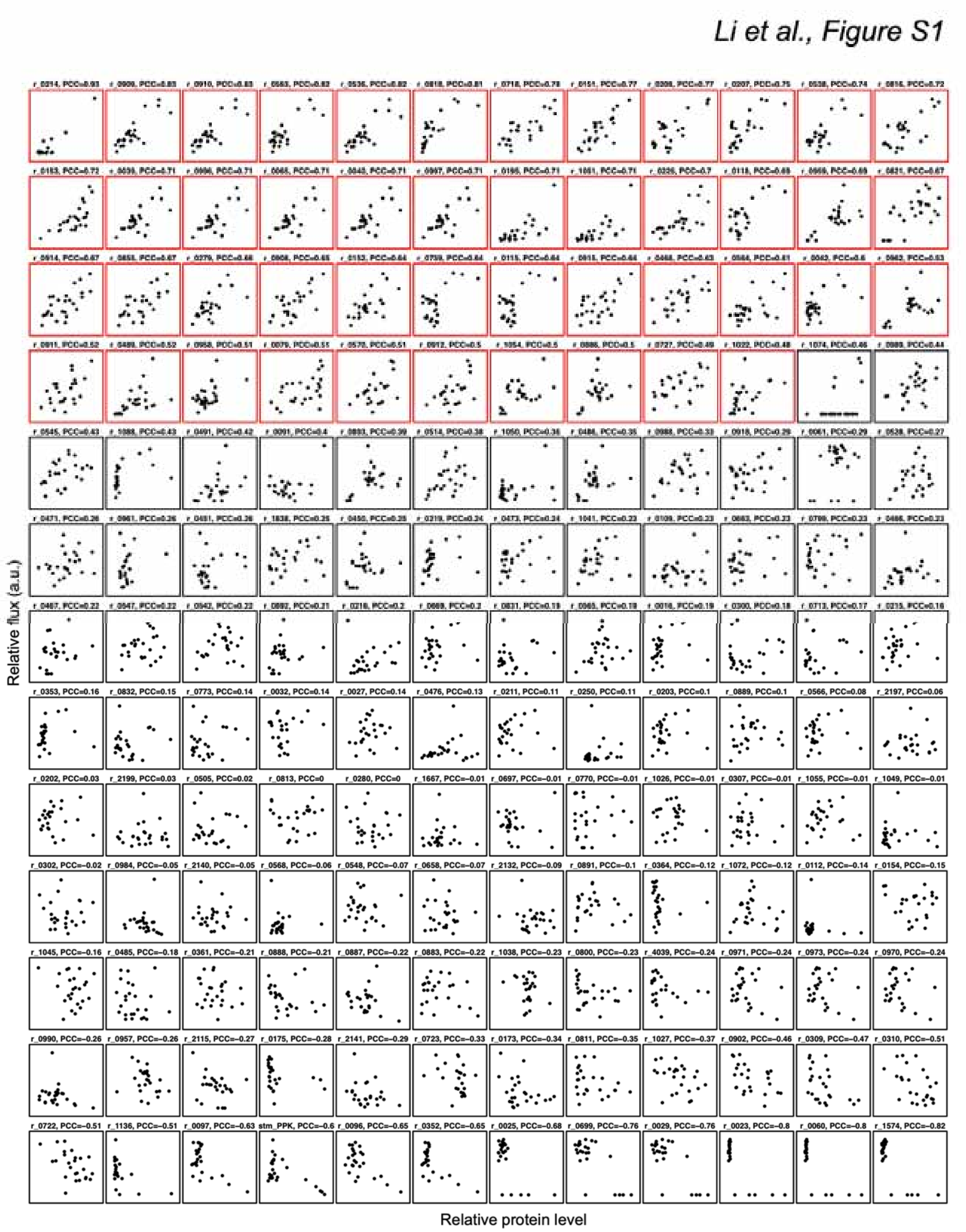


**Figure S1: Scatter plots showing the flux-enzyme level correlation for all 156 testable reactions in yeast.**

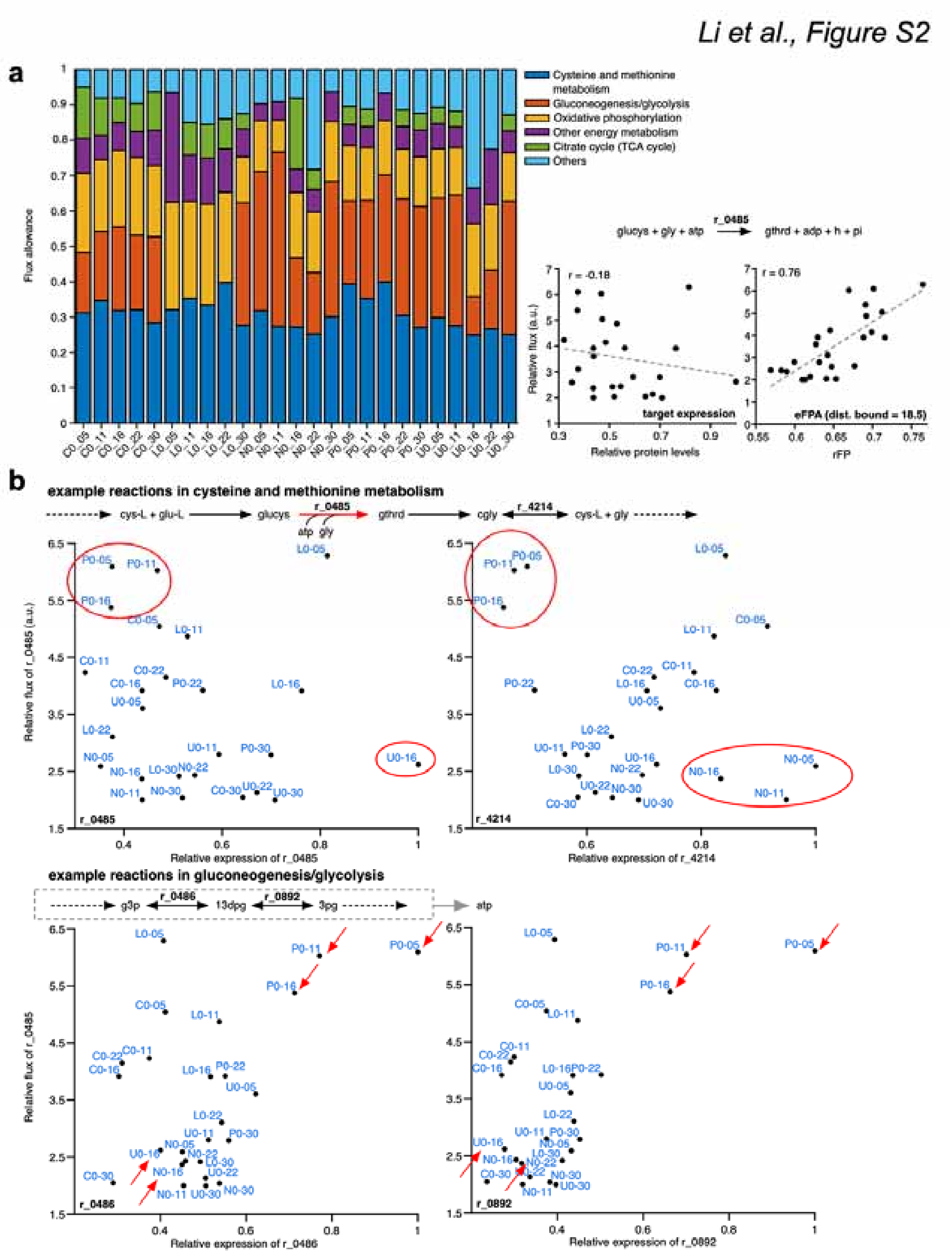


**Figure S2: Mechanisms for the prediction of glutathione synthesis flux a**, Left panel: flux allowance contributions of different metabolic pathways (colors) in different conditions (x-axis) during the optimal-boundary eFPA of the glutathione synthesis reaction r_0485, which belongs to the *cysteine and methionine metabolism* in the pathway annotations. Flux allowance contribution is defined by the product of reaction weight coefficient and flux (see Supplementary Methods for details). Since total allowance is held constant, major contributors to this value have the biggest influence on the prediction of the relative flux of the target. Right panel: scatter plot showing the correlation between rFP and relative flux of glutathione synthesis. **b**, Scatter plot showing the correlation between the relative protein levels of key contributing reactions and the flux of glutathione synthesis. The red circles or arrows are used to highlight in all graphs the conditions in which the enzyme levels of glycolysis reactions are concordant with the flux of glutathione synthesis while that of reactions in close proximity to the target are not.

*Li et al., Figure S3*

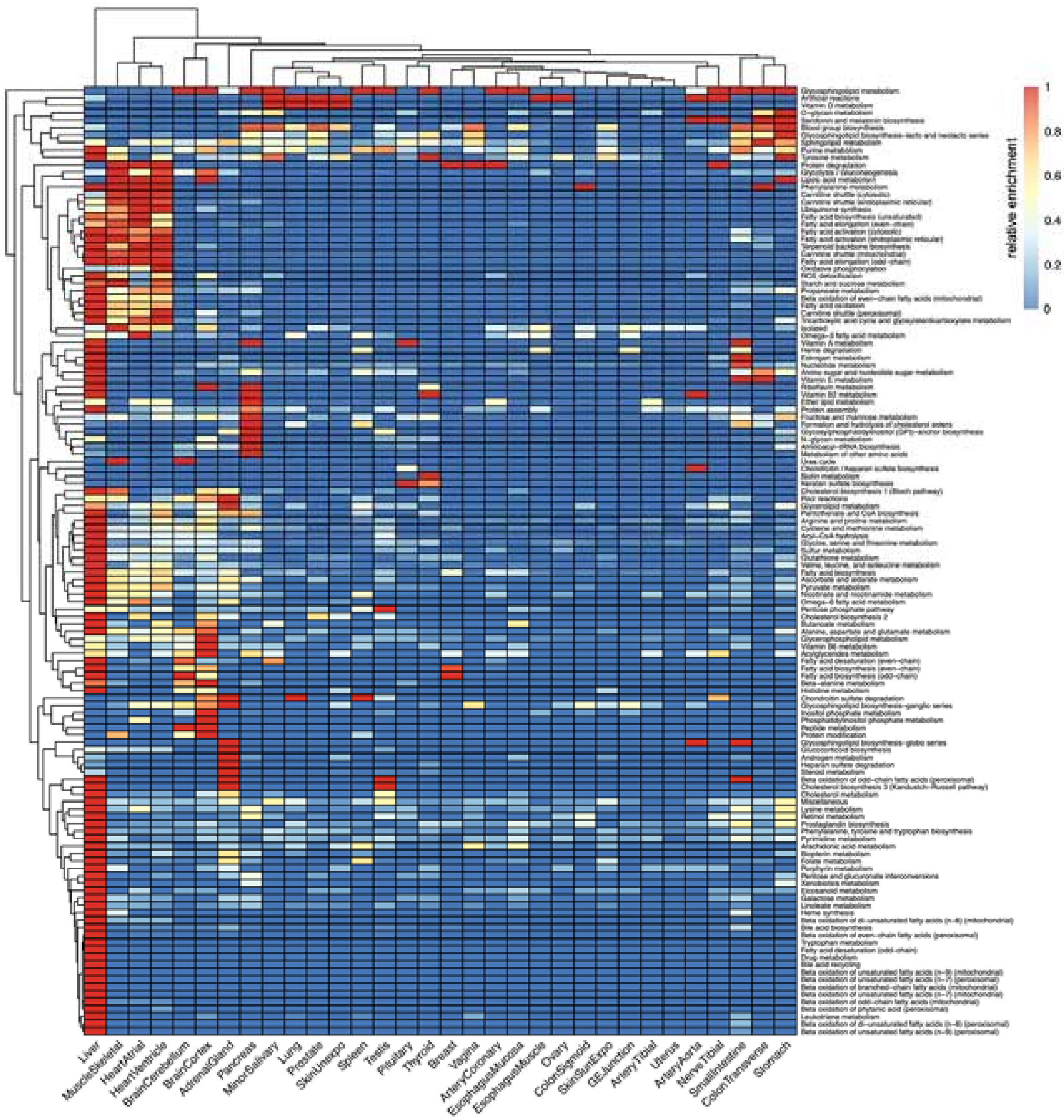


**Figure S3: Heatmap showing the relative tissue-enrichment of different**

**subsystems in human 1 model.** The levels of relative enrichment for indicated subsystems (rows) in each human tissue analyzed (columns) are shown. A reaction is considered enriched in a tissue if its tissue enrichment score (ΔrFP) in that tissue is greater than 0.2. The number of tissue-enriched reactions for a subsystem and tissue is row-wise (across tissues) normalized by dividing with the row maximum to derive the relative enrichment. This heatmap is based on the eFPA predictions with the protein data.

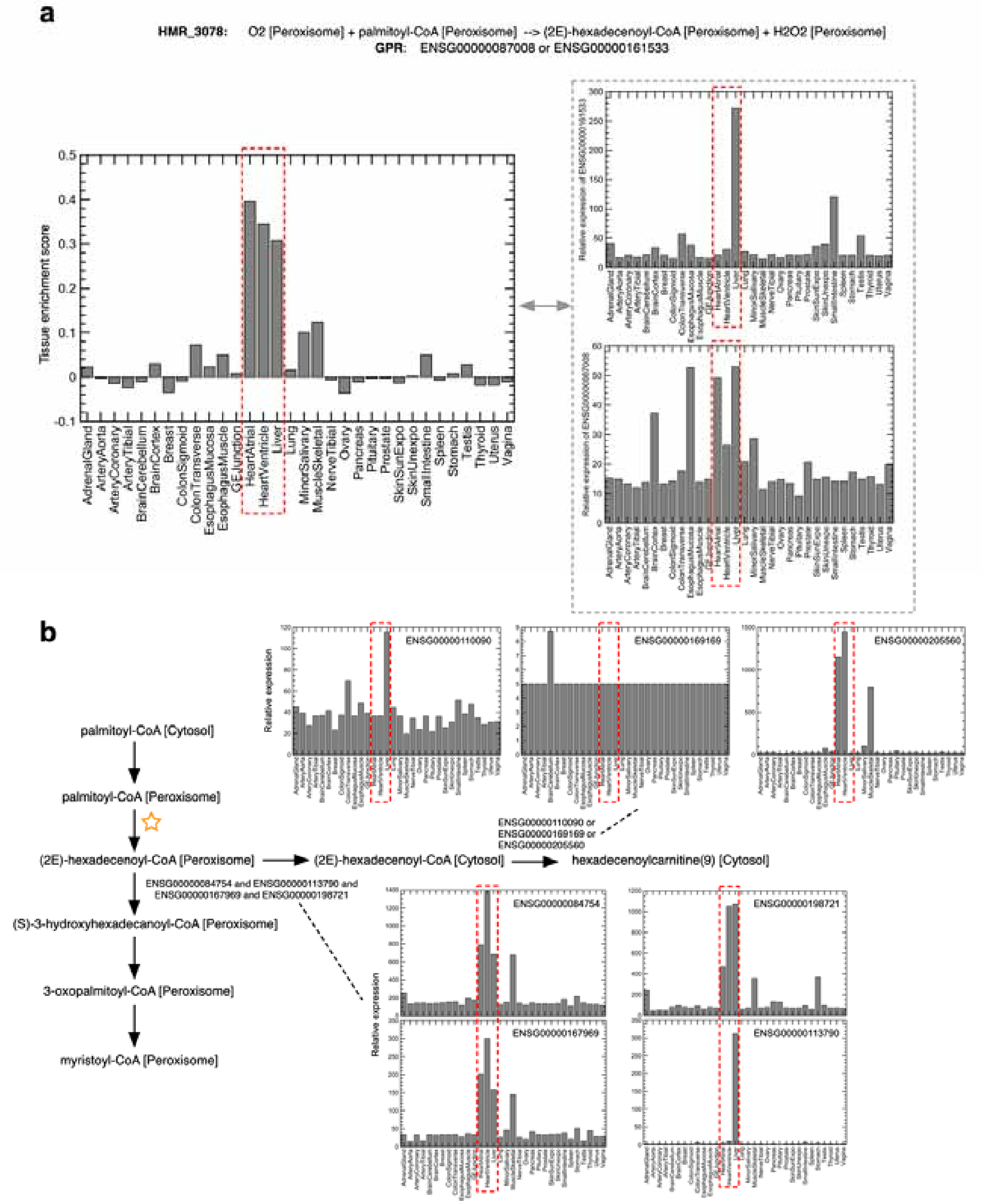


**Figure S4: An example of eFPA-driven prediction of human tissue metabolism. a**,

ΔrFP prediction for palmitoyl-CoA peroxisomal oxidation. The relative protein levels of two associated enzymes are plotted on the right. Although liver expression was consistently high in both enzymes, the heart expression was not. **b**, Enzyme levels of surrounding reactions for palmitoyl-CoA peroxisomal oxidation. The product of the target reaction is either further oxidized in peroxisome or exported into cytosol. The gene expressions in both routes show significant enrichment in heart.
